# Design, infectability, and transcriptomic analysis of transregionally differentiated and scalable lung organoids derived from adult bronchial cells

**DOI:** 10.1101/2024.07.02.601655

**Authors:** Alicia Reyes Valenzuela, Mark Turner, Nathan Markarian, Christophe Lachance-Brais, John Hanrahan, Hojatollah Vali, Silvia Vidal, Luc Mongeau

## Abstract

The lung is a primary target for many lethal respiratory viruses, leading to significant global mortality. Current organoid models fail to completely mimic the cellular diversity and intricate tubular and branching structures of the human lung. Lung organoids derived from adult primary cells have so far only included cells from the input cell region, proximal or distal. Existing models are expensive. They often require cells from invasive deep lung tissue biopsies. The present study aimed to address these limitations. The lung organoids obtained using an original protocol exhibited transregional differentiation and were derived from relatively more accessible primary cells from the trachea/bronchi. Immortal bronchial cell lines were also used to simplify organoid fabrication and improve its scalability. The lung organoids are formed starting from bronchial cells with fibroblasts feeder cells in an alginate hydrogel coated with base membrane zone proteins. Characterizations were performed using bulk RNA sequencing and tandem mass tags. The resulting organoids express markers of different lung regions and mimic to some extent the tubular and branching morphology of the lung. The proteomic profile of organoid from primary cells and from cell lines was found to evolve towards that of mature lung tissue. Upregulated genes were mostly related to the respiratory system, tube development, and various aspects of respiratory viral infections. Infection with SARS-CoV-2 and influenza H1N1 was successful and did not require organoid disassembly. The organoids matured within 21 days and did not require complex or expensive culture methods. Transregionally differentiated lung organoid may find applications for the study of emerging or re-emerging viral infections and fostering the development of novel in-vitro therapeutic strategies.

## INTRODUCTION

The human lung is the primary target for many lethal respiratory viruses. Influenza viruses and coronaviruses, for example, can cause acute respiratory distress syndrome^1–3^. Lower respiratory infections constitute the fourth leading cause of death worldwide^4^. Viral infections depend on host-specific mechanisms. Given their rapid adaptation to the host environment, the circulating viruses can be genetically distinct from those grown under standard laboratory conditions such as 2D monocultures^3^. These genetic alterations typically occur in virus envelope proteins that are crucial for viral entry and transmission^3^. This is the case for the human parainfluenza virus^3^, but it could potentially be true for other viruses as well.

It is generally acknowledged that the use of animal models for viral infections may be misleading due to inter-species biological and individual differences, as well as interference with other microorganisms^5^. Conventional 2D monocultures are also problematic, as they do not mimic tissue organization *in vivo*, nor cell integrity due to 2D culture adaptation^5^.

Organoids, in contrast, can better recapitulate the cellular organization and physiology of original organs, including cell communication^6^. Physiological mimicry is particularly useful, if not essential, for the study of viral-host interactions, which involve more than one cell type in the targeted tissue. For example, severe acute respiratory syndrome-2, SARS-CoV-2, first infects cells from the proximal lung such as airway epithelial cells, ciliated and goblet cells. If not cleared, the virus later infects cells from the distal lung region such as alveolar type II (AT2) cells as its primary target^7^. Influenza A, as another example, infects airway epithelium cells lining the lung, including AT2, thereby destroying key gas-exchange mediators and allowing viral exposure to endothelial cells^8^. Gut, brain, vascular, kidney, and lung organoids have been used to model SARS-CoV-2 and other infectious diseases^1^. Organoids have yielded improved predictive results for drug testing before clinical trials. Their use is increasingly being adopted by pharmaceutical companies^9^.

Because of their prospective value as viral infection models, there has been a need for physiologically multi-regional lung organoids. So far, organoids specifically designed to model the lung have remained limited in their intrinsic complexity^4^. Most existing organoids target one specific region of the lung, either proximal or distal. Examples include alveospheres^10^, broncheospheres^11^, tracheospheres^12,13^ or human airways organoids^14^. As shown in Table S1, existing organoids have been generated by primary cells derived from deep lung human or mice tissue embedded in Matrigel. Recent progress has been made towards more intricate lung organoid models. Zhou, and colleagues developed protocols for the generation of independent proximal and distal lung organoids. But these regional models are independent, hampering the study of viral infection with viral tropism as in the human body^15^. Das and colleagues created a lung organoid expressing markers of both proximal and distal regions by modifying the cell culture medium with additional growth factors and antioxidants^16^.

The scalability of lung organoids production remains problematic. Except of organoids derived by induced pluripotent stem cells (iPSC), lung organoid models require deep lung tissue biopsies for the harvesting of primary cells needed for their fabrication^1,10–18^. Cancer and other diseased cells, are often used in lieu of healthy cells, given the lack of availability of healthy donors, and the invasive nature, and costs of lung tissue biopsy^19,20^. Clever and colleagues established culture conditions for the long-term expansion of airway organoids for up to one year. But these lung organoids only include cell populations from proximal regions such as basal, secretory and ciliated cells; they do not express distal lung markers^21^. One common characteristic of the lung organoids derived from primary adult cells is their spheroidal morphology (S. Table 1). Spheroids do not reproduce the complex branching structures associated with the lung^1,10–18^. More complex lung organoids derived from induced pluripotent stem cells have recapitulated multiple lung regions, as well as a more arborized morphology^17,18,22,23^, as shown in S. Table 1. But, these organoids require around 85-150 days to grow, along with costly and complex culture conditions^17^. Such growth rate is insufficient to track epidemic progression and viral evolution in a crisis. In addition, they display immature phenotypes, and thus cannot be infected with adult viral diseases, such as influenza type A and SARS-CoV-2^22^. Some lung organoids derived from primary adult cells have shown infectibility with SARS-CoV-2 and influenza; but the need for segmentation and culture in cellular monolayers^14,16,19^ negates the potential benefit of using organoids to understand disease progression.

Organoid research has, so far, been confronted with the use of physiologically accurate models, which are complex and multi-regional but cannot be infected due to their immature phenotype^17,18,22,23^, or mature models that are infectible but only represent one specific region of the lung with limited complexity^14,16,19^.

The objective of the present study was to create lung organoids featuring transregional differentiation, defined as the differentiation of the input cells into phenotypes from both the proximal and distal regions of the lung (Fig. 1A). Additionally, the study aimed to achieve several desirable characteristics for the organoids: a complex tubular morphology indicative of self-organization, low cost, scalability, and susceptibility to infection. Primary epithelial cells from the trachea/bronchi, within the proximal lung region, were used as progenitors. The protocol led to their differentiation into phenotypes from the distal regions of the lung, while maintaining some phenotype characteristics from the proximal region. The same protocol was replicated using immortal bronchial epithelial cells to further reduce model costs and enhance its availability. One more aim was to infect these organoids with SARS-CoV-2 and influenza H1N1 without the need of their segmentation to prevent viral 2D culture adaptions ^24^.

**Fig. 1.**
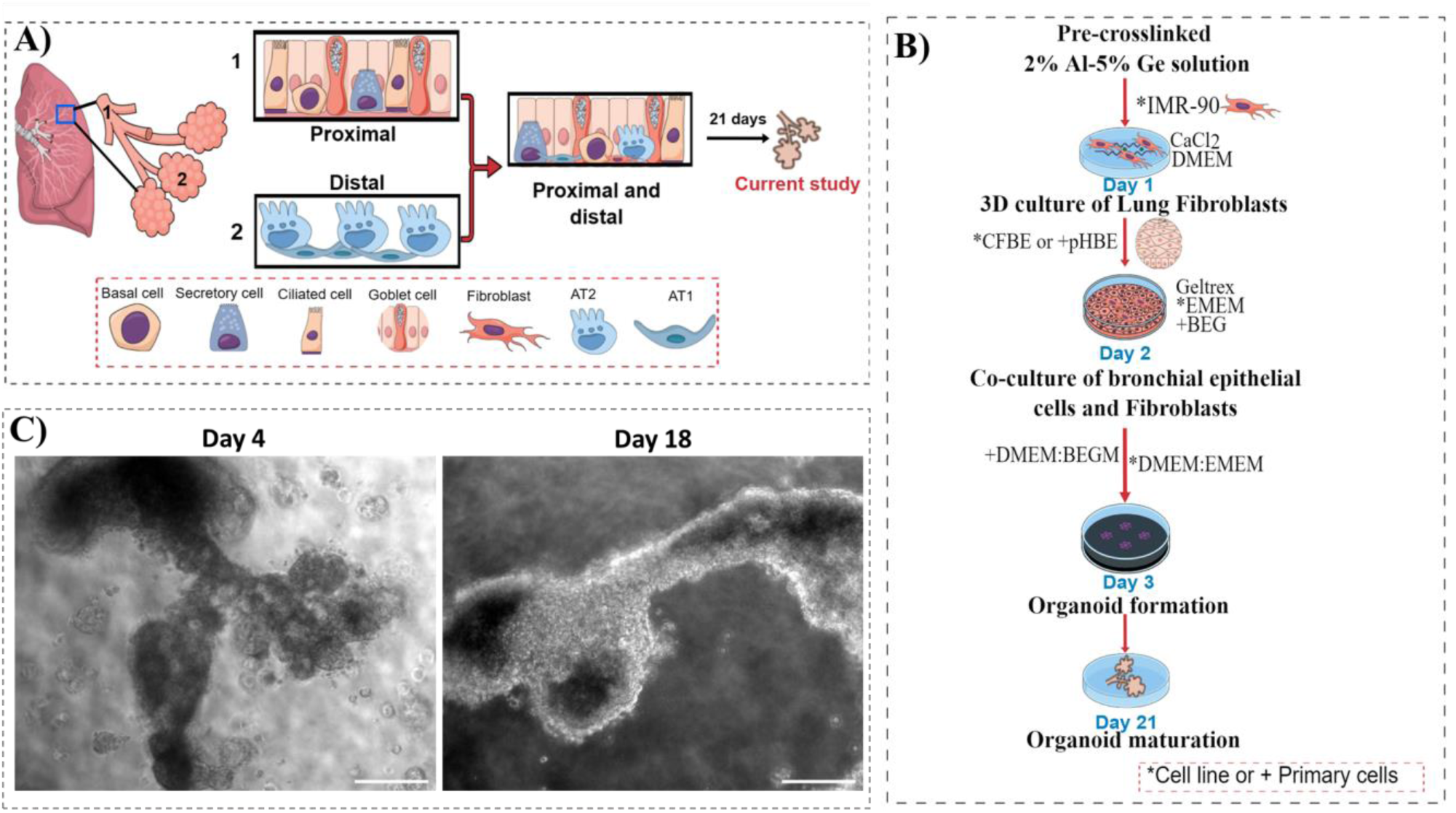
Generation of lung organoids. A) Simplified schematic of the lung, as composed of two distinctive regions: proximal and distal. Transregional lung organoids contain cells from both regions. B) Protocol for the development of lung organoids from the co-culture of lung fibroblasts and airway epithelial cells from primary or immortalized origin. C) Representative phase contrast pictures of lung organoids derived from immortal cell lines. Magnification 10X. Scale bar 200 um.

The following results describe lung organoids encompassing markers of cell types from both the proximal and distal regions of the lung; these are referred as transregional differentiated lung organoids (TDLO). These organoids could be formed from primary bronchial cells obtained from patients (TDLO-p). A more scalable version of lung organoids (SLO-i) was achieved by using immortalized cell lines. Both types of organoids display a tubular and branching morphology. The maturation of TDLO-p and SLO-i was reached within a 21-day period. As a proof of concept, SLO-i was successfully infected using SARS-CoV-2 and influenza H1N1. Viral replication was quantified over time. These new models combine lung mimicry and facile preparation, to possibly improve upon existing organoid models for viral drug testing.

## RESULTS

### Creation of human lung organoids derived from primary or immortalized bronchial epithelial cells

To induce the formation of human lung organoids with a mimetic morphology, fibroblasts were co-cultured with airway epithelial cells. The interaction between these cell types was to allow the secretion of endogenous growth factors by the fibroblasts and guide organoid development. This approach differs from the typical monoculture of airway epithelial cells with exogenous fibroblast growth factors^10,11,13–16,21^. Epithelial progenitors, surrounded by mesenchyme, trigger the development of native lungs^25,26^. To emulate this process, cells from a fibroblast finite cell line (IMR-90), were encapsulated into a thin layer of hydrogels consisting of 2% Alginate and 5% Gelatin (AlGe). This specific substrate composition was chosen for its initial stiffness of 2.025 kPa, which is representative of that of the lung and other soft tissues^27^. On day one, cells were cultured with Dulbecco’s Modified Eagle Medium (DMEM). The following day, base membrane zone proteins such as laminin, collagen IV, entactin, and heparin sulfate proteoglycans (Geltrex) were incorporated into the superior portion of the hydrogel forming a secondary thin layer. Primary human bronchial epithelial cells, HBE (isolated from human bronchus/trachea of both sexes, namely one 53-year-old female and one 51-year-old male) were then seeded on the top of the secondary layer.

These primary cells were cultured with Bronchial Epithelial Cell Growth Medium (BEGM). The input cell population contains basal cells, positive for p63^14,15,28^. It was been found that these basal cells differentiate into multiple types of airway epithelial cells, including the ones populating the distal lung regions^29^. This process can occur following lung injury, as reported in preclinical and clinical studies^29^. Fibroblasts growth factors are known for their important roles in lung development^25,26^, in particular the proliferation and stemness maintenance of stem cells ^12,30,31^. We hypothesized that fibroblast cues would induce basal cell differentiation into distal regions of the lung, including AT2 and AT1 cells. At day 3 of culture, cells were maintained in medium combining of BEGM: DMEM in 1:1 ratio until day 21. The cell medium was changed every two days. An overview of the protocol is shown in Fig. 1B.

The same approach was used with cell lines, parental human cystic fibrosis bronchial epithelial cells were used, CFBE41-o. Although this cell line was originally derived from patients with cystic fibrosis expressing F508 Del mutation^32^, these cells are devoid of any CFTR expression once immortalized^32,33^. They are convenient, low-cost source of bronchial epithelial cells given their commercial availability. It is well known that during the immortalization process of primary and stem cells, there is a potential loss of their differentiation potential^34^. It has been suggested that the culture of these cells into a proper microenvironment may help to overcome their loss of stemness ^34–38^. In a few studies, the culture of immortalized cell lines has led to their differentiation^37,38^. We hypothesized that our protocol would trigger the differentiation of CFBE41-o into organoids with proximal and distal lung phenotypes.

We monitored the development of lung organoids generated either with primary cells (LOp) or immortalized cells (LOi) using low phase contrast microscopy. Figure 1.C shows representative pictures of the derived LOi. Their morphology is not spheroidal but features tubular-like elongated structures growing in thickness and diameter over time to reach an average size of 1.07 ± 0.06 cm at day 21.

### Cellular population and functional enrichment analysis for lung organoids derived from primary cells

The objective was to identify the cellular differentiation of primary cells into the six major lung epithelial cells from proximal and distal region^14–16,21,39^ of LOp, using bulk RNA sequencing. Other markers for rare cell types were also evaluated^26,39,40^. Additionally, the morphology of these organoids was monitored over time.

Nine organoids were processed and divided into three samples, each consisting of three organoids (n=3 per sample). This grouping was chosen to ensure an adequate amount of RNA for analysis. We compared the organoid’s gene expression with the one from the input cells: primary bronchial epithelial cells, HBE, cultured in filters coated with collagen IV (see methods details), creating an air-liquid interface, ALI, (n=3). This model is well-known to be the gold standard for proximal airways^41^.

Similarly, the organoid gene expression was compared with the one from fibroblasts cultured in traditional monolayers (n=3).

We used multiple markers for different cell types, as illustrated in Fig.2A. These markers have been extensively used before for the identification of specific cell types^10,14–17,21,23,39^, since they are the most abundant cells in the lung^26^. In addition, we used SOX2 as marker of multipotency of proximal region of the lung^22,26,42^, while SOX9 was used as a marker of multipotency of distal region of the lung^22,26^. For rare and recently discovered cells, we aimed to identify markers associated with brush cells^26^, and pulmonary neuroendocrinal cells (PNES)^26^, located on the proximal lung regions.

**Fig. 2.**
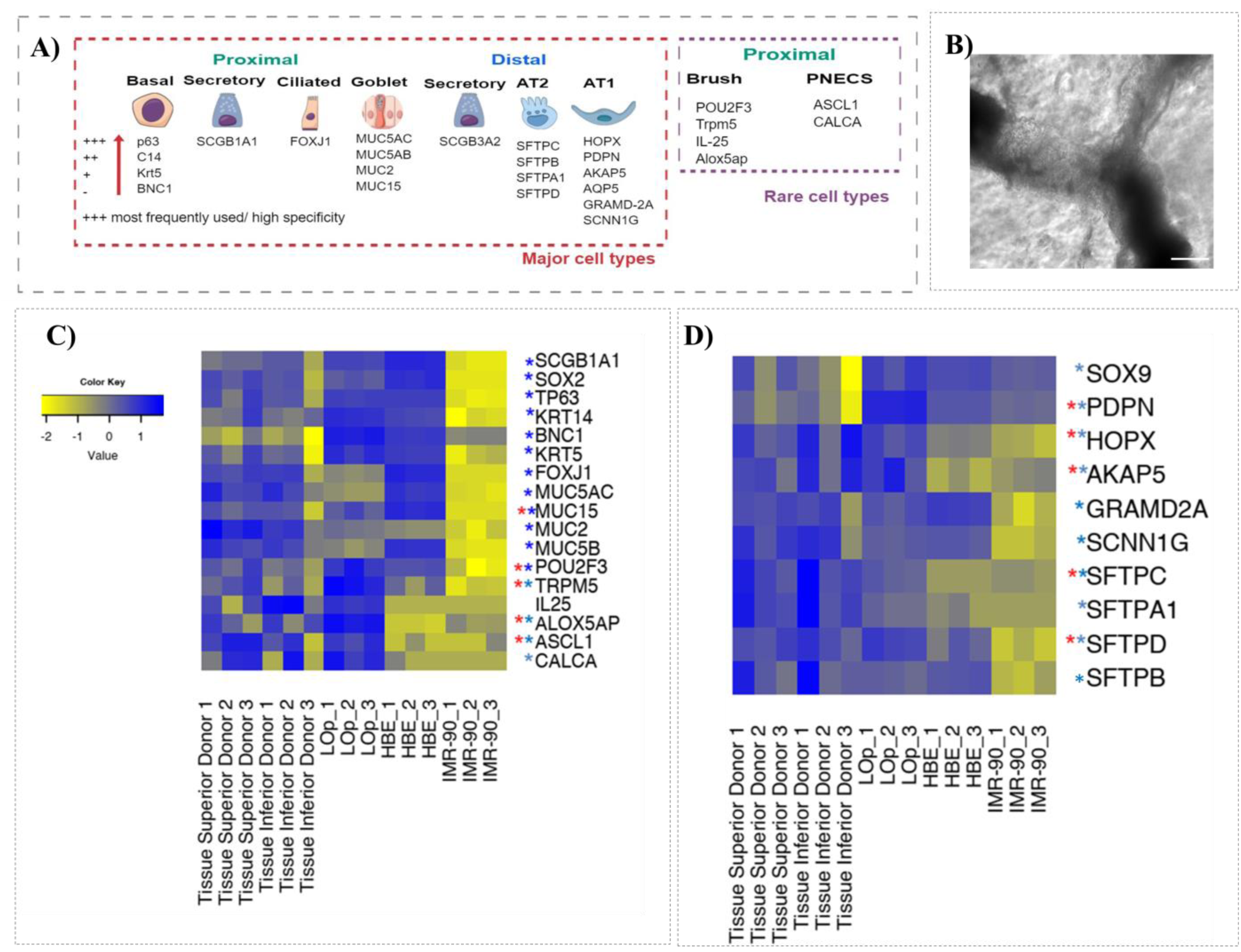
Characterization of lung organoids derived from primary bronchial epithelial cells (LOp) A). Markers used for the identification of major or rare cell types in the lung in either proximal or distal regions. B). Representative phase contrast picture of female derived lung *organoid* at day 18. Magnification 10x. Scale Bar 200 um. C-D). RNA sequencing data showing proximal and distal markers of LOp, fibroblasts monolayers, and primary bronchial epithelial cells, respectively, in air-liquid-interface. N=3. Differentially expressed genes (DEGs) were defined using adj-pvalue < X and |log2FCshrink|≥4 X leading respectively to X, Y, Z DEGs for A vs B, A vs C and B vs C. DEGs. * p.adjust <0.05 in comparison to IMR-90, *p.adjust <0.05 in comparison to HBE.

Lung organoids derived from primary cells show a tubular morphology with some ramifications (Fig. 2B).

The expression from the markers used to identify basal cells, shown in Fig.2C, Cytokeratin 14, Keratin 5, BNC1, p63, remains the same between HBE monolayers (grown in ALI) and TDLO-p. As expected, monolayers of IMR-90 were devoid of any marker of basal cells, nor any other cell types from the proximal lung region. It was found that the expression of markers associated with secretory cells (SCGB1A1+), ciliated cells (FOXJ1+), and multipotency (SOX2+) decreased within the organoid in comparison to HBE monolayers. Yet their expression was significantly greater than that in IMR-90 cell monolayers. It was found that markers of goblet cells (MUC5AC, MUC5B, and MUC2) were upregulated within HBE cells in comparison to LOp. Only mucin 15 was found to be upregulated in LOp. Most of the cell markers used to identify brush cells (POU2F3, TRMP5, ALOX5AP), and PNECs (ASCL1) were found to be upregulated within the organoid in comparison to HBE and IMR-90 (Fig. 2C).

Markers of cells from distal regions of the lung such as secretory (SCGB3A2+), Alveolar type II (SFTPC+, SFTPD+) and Alveolar type I (HOPX+, PDPN+, AKAP5+) were all upregulated within the organoid in comparison to HBE and IMR-90 cell monolayers as shown in Fig. 2C. SFTPB and SFTPA were also present on HBE cell monolayers.

Overall, markers associated with proximal regions of the lung were down regulated in the organoid, while markers associated with distal regions of the lung were upregulated. These observations suggest that input bronchial cells, HBE, from proximal regions, differentiate into multiple cell types from distal region while retaining some markers of the proximal region. Because of this feature, these organoids were named Transregionally Differentiated Lung Organoids (TDLO-p). The expression of representative genes of alveolar type II cells (SFTPC+) is also supported by flow cytometry results. These results indicate that at least 10% ± 3% of the cells conforming the organoid are positive for SFTPC (S. Fig.1).

A quantitative comparison between gene expression in cell monolayers used to form the organoids (HBE and IMR-90) and LOp (p adj.<0.05) revealed that 153 genes were up- or downregulated. A functional enrichment analysis of those genes was performed using g: Profiler. Gene Ontology (GO) analysis identified genes from several domains: Molecular function (MF), Biological Process (BP), and cellular components (CC), as well as other biological databases as shown in S.Fig.2. Fig. 2 shows multiple BP terms identified from the functional enrichment analysis including respiratory system development, tube development and lung alveolus development. Many genes from each term were upregulated in the organoid in comparison to cell monolayers (Fig. 2A-C).

These results support the finding that the organoid constitutes a more transregionally differentiated respiratory system model compared to HBE or fibroblasts alone. In addition, genes associated with tube development (Fig. 2B) were upregulated within the organoid, which suggests that the tubular morphology of the organoids may be the consequence of well orchestrated and complex processes. This is also supported by other related terms, such as branching morphogenesis, tube morphogenesis, and morphogenesis of branching structure. Once organoids are formed, genes from each individual cell type, HBE or IMR-90 are upregulated. Additional genes are original and not abundantly present in the input cells. This shows that the organoids evolve and self-organize to yield a model that is more than the sum of the characteristics of the input of cells.

### Cellular population of lung organoids derived from immortalized cell lines

We aimed to evaluate the protocol for lung organoids using immortalized cell lines (LOi) instead of primary epithelial cells. Parental human cystic fibrosis bronchial epithelial cells CFBE41-o. were used. It was observed that the elimination of fibroblasts from the protocol led to a spheroidal morphology (Fig. 4A). The addition of fibroblasts to the LOi led to a branching tubular morphology (Fig. 4B). This supported our hypothesis that fibroblasts are critical for obtaining a more sophisticated morphology.

**Fig. 3.**
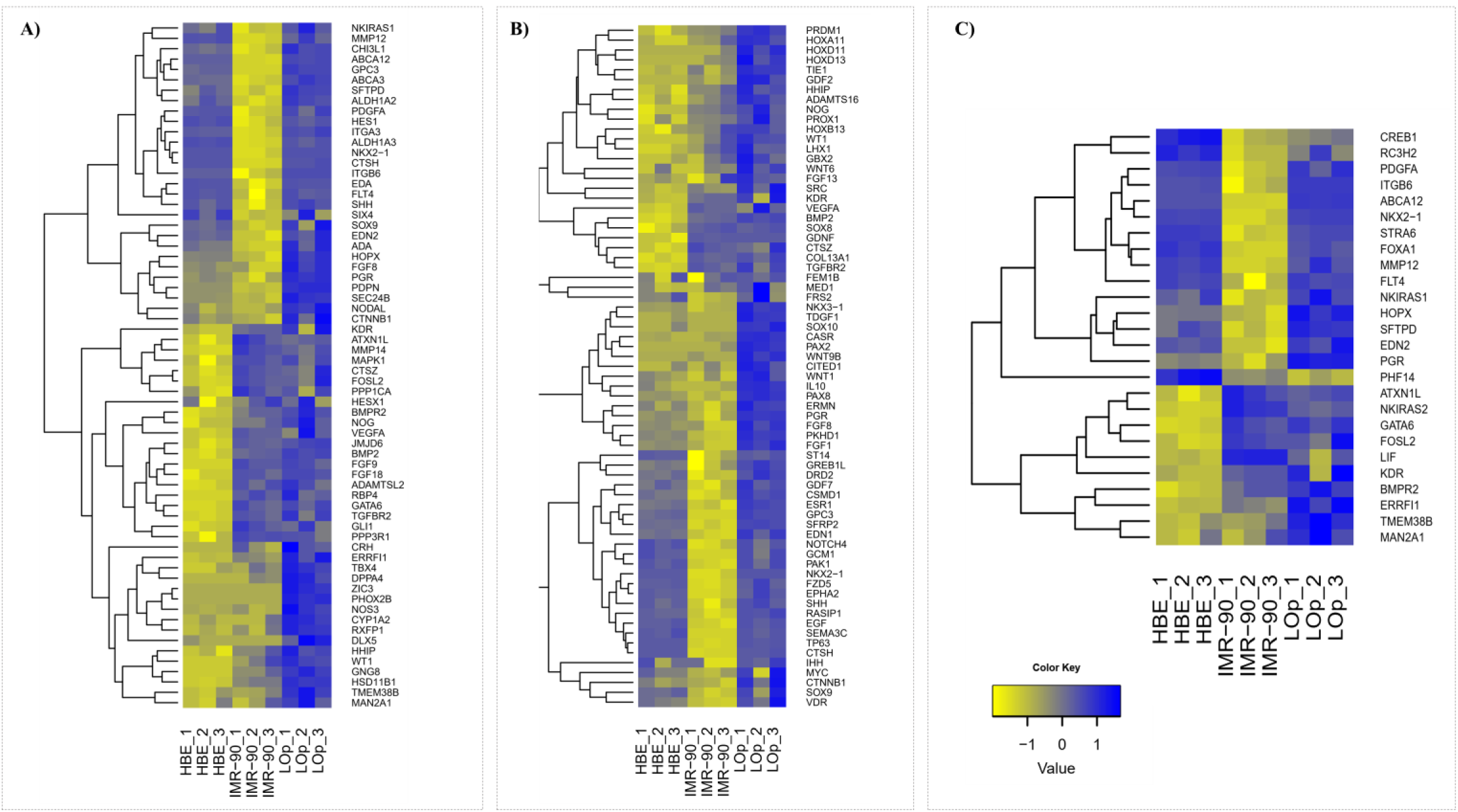
Functional Enrichment Analysis of lung organoids derived from primary cells (LOp), primary epithelial cells (HBE) and lung fibroblasts (IMR-90) using g: Profiler. N=3. A-C) Heatmaps showing genes associated with respiratory system development (A), tube development (B), and lung alveolus development (C) in all samples. Upregulation shown in blue. Differentially expressed genes (DEGs) were defined using adj-pvalue < X and |log2FCshrink|≥4 X leading respectively to X, Y, Z DEGs for A vs B, A vs C and B vs C. DEGs.

**Fig. 4.**
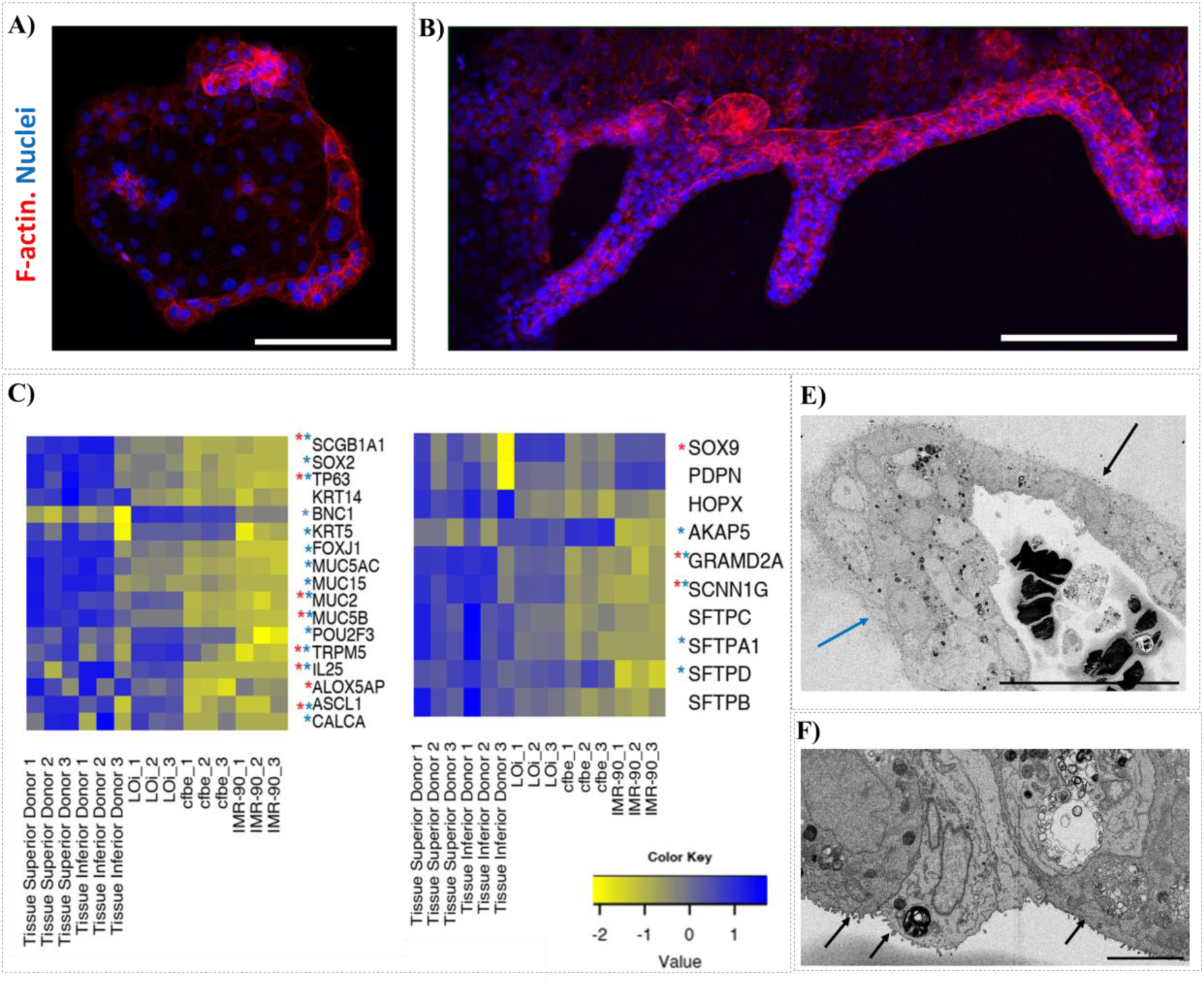
Characterization of lung organoids derived from immortal bronchial epithelial cells (LOi). A) Images showing the morphology of organoids without fibroblasts in their culture at day 7. B) Images showing typical tubular morphology of lung organoids at day 7. Red-F-actin. Blue-Nuclei. Magnification 10X. Scale Bar 200 µm. C-D) N=3. Differentially expressed genes (DEGs) were defined using adj-pvalue < X and |log2FCshrink|≥4 X leading respectively X, Y, Z DEGs for A vs B, A vs C and B vs C. DEGs. * p.adjust <0.05 in comparison to IMR-90, *p.adjust <0.05 in comparison to CFBE. E-F) Organoids (LOi) at day 21 were visualized using Focused Ion Bean microscopy (FIB). E) Structure of LOi. Pseudostratified squamous epithelium (black arrow) and stratified squamous epithelium (blue arrow) were identified within the organoid. Magnification 1076X, scale bar 50 µm. F) Microvilli structures (black arrows) and signs of autophagy were found in LOi. Magnification 5000X. scale bar 5 µm.

The transcriptome of LOi was studied using RNA-sequencing using the same markers as for LOp (Fig. 2A). Low levels of markers of basal cells (Cytokeratin 14, and p63) were found in the initial cell monolayers (Fig. 4C), as expected given their stage of immortalization. After organoid formation, p63 was upregulated in comparison to CFBE and IMR-90 (Fig. 4C). This indicated that the protocol triggered changes in the genome even after cells are immortalized^37,43^, presumably driving by microenvironment.

Markers of secretory cells (SGB1A1+) from the proximal region, goblet cells (MUC5B+, MUC2+), brush cells (TRMP5+, IL25+), and PNECS cells (ASCL1+), were all undetectable or minimal in cell monolayers, but they were all significatively upregulated in LOi (Fig.4C). Similarly, secretory cells (SGC3A2+) from distal lung regions were upregulated in the organoids. Alveolar type II markers, and AT1 markers were found not to be upregulated in the LOi in comparison to cell monolayers. Nevertheless, the transcriptome profile of the LOi is more similar to the one of adult tissue biopsies (male donors aged 42, 51, and 66 years) than to the transcriptome of the input cell lines used to form the organoids (Fig. 4C-D). Altogether, gene expression data indicates changes in LOi, regardless of the initial immortalization stage of the cells. But no cell differentiation into distal phenotypes was observed. Therefore, TDLO could not be achieved using immortal cell lines. The expression of various markers used was less abundant than that in organoids derived from primary cells. Nevertheless, the expression was significantly greater than that in cell monocultures.

Focused Ion Bean (FIB) microscopy was used to evaluate the morphology of LOi at day 21. Figure 4E shows LOi with a magnification of 1076 X. Arrows indicate the main organoid structure, including morphological heterogenicity. Blue arrows point at multiple layers of epithelial cells, while black arrows shows a pseudostratified squamous epithelium, characteristic of the airways of the respiratory system^44^. Cell debris was observed near the center of the organoid. These observations underscore the complexity and heterogeneity of cellular self-organization within the organoid and shed light on its structural diversity. With a magnification of 5000 X (Fig.4F), microvilli structures were identified, as shown in Fig.4F. Microvilli are microscopic membrane cellular protrusions which are involved in the process of absorption of nutrients and cellular secretion^45^. The emergence of microvilli suggests a level of cellular differentiation and specialization within the organoids, despite their origin from cell lines that are traditionally considered homogeneous^41,46^. This finding challenges the notion of cellular homogeneity, within the organoids and underscores their dynamic nature, potentially reflecting a degree of cellular plasticity and functional diversity. We anticipate observing similar structural heterogeneity for TDLO-p. Notably, the identification of microvilli, which is typically associated with brush cells in the lung^47^, further highlights the organoids’ ability to mimic essential characteristics of native lung tissue. Evidence of autophagy was discerned in LOi (Fig. 4F). Autophagy is the process of degrading cytoplasmic components within lysosomes^48^. This process serves various physiological functions, including adaptation to starvation, clearance of intracellular proteins and organelles, development, anti-aging, elimination of microorganisms, cell death, tumor suppression, and antigen presentation^48^. Interestingly, autophagy has been associated with the regulation of lamellar body formation^49^. Lamellar bodies are concentric structures loaded with surfactant proteins specific to AT2 cells^49^. While the reason for the abundance of autophagy in the organoids is not known, one possibility could be its involvement in the development of these organelles.

### Proteomic profiling of organoids and cell lines

Additional indications of cellular dynamics were obtained using Tandem Mass Tags (TMT). Three samples (n=3) of each type were used for this analysis. A principal component analysis of the entire dataset (including lung tissues), depicted in Fig. 5A, indicated that both types of organoids, LOp and LOi are reproducible, as evidenced by the tight clustering of their respective samples (Fig.5A). A detailed numerical analysis of proteins found in the input cell monolayers HBE, IMR-90, and LOp, is shown in Figure 5B. Among these samples, 1,361 proteins were shared. Specifically, 1,542 proteins were identified in LOp, of which 180 were exclusive to the organoid and absent from the cell monolayers. Similarly, a numerical comparison of proteins expressed in CFBE, IMR-90, and LOi is shown in Figure 5C. A total of 1,357 proteins were common to these three groups. In the LOi, 1,542 proteins were identified, with 180 proteins exclusively found in the organoids. A closer look of the upregulated proteins in LOp and LOi is shown in Fig. 5D and Fig.5E, respectively, as well as in S.Fig.3-4, and S. Table 2-3. Surfactant protein A stands out among the upregulated proteins of the LOi. This protein plays an important role in the host-defense mechanisms of the lung, including opsonizing virus, bacteria, and apoptotic cells; it also modulates the production of cytokines and mediators of the inflammatory response^50^. In organoids derived from primary cells, Prolactin Induced Protein (PIP) stands out. According to the human protein atlas, this protein is characteristic of respiratory ciliated cells. All the upregulated proteins in the organoids were analyzed using STRING software: https://string-db.org/ to collect supporting information from multiple sources such as text mining of scientific literature, databases of interaction experiments, and complex pathways^51^. All proteins were associated with lung tissue. It was also observed that these proteins formed interconnected networks. This facilitates coordinated communication and regulation of essential cellular functions, as shown in S.Fig.5. Protein networks enhance signal amplification, pathway regulation, and adaptation to changing cellular conditions compared to proteins working in isolation^52^.

**Fig. 5.**
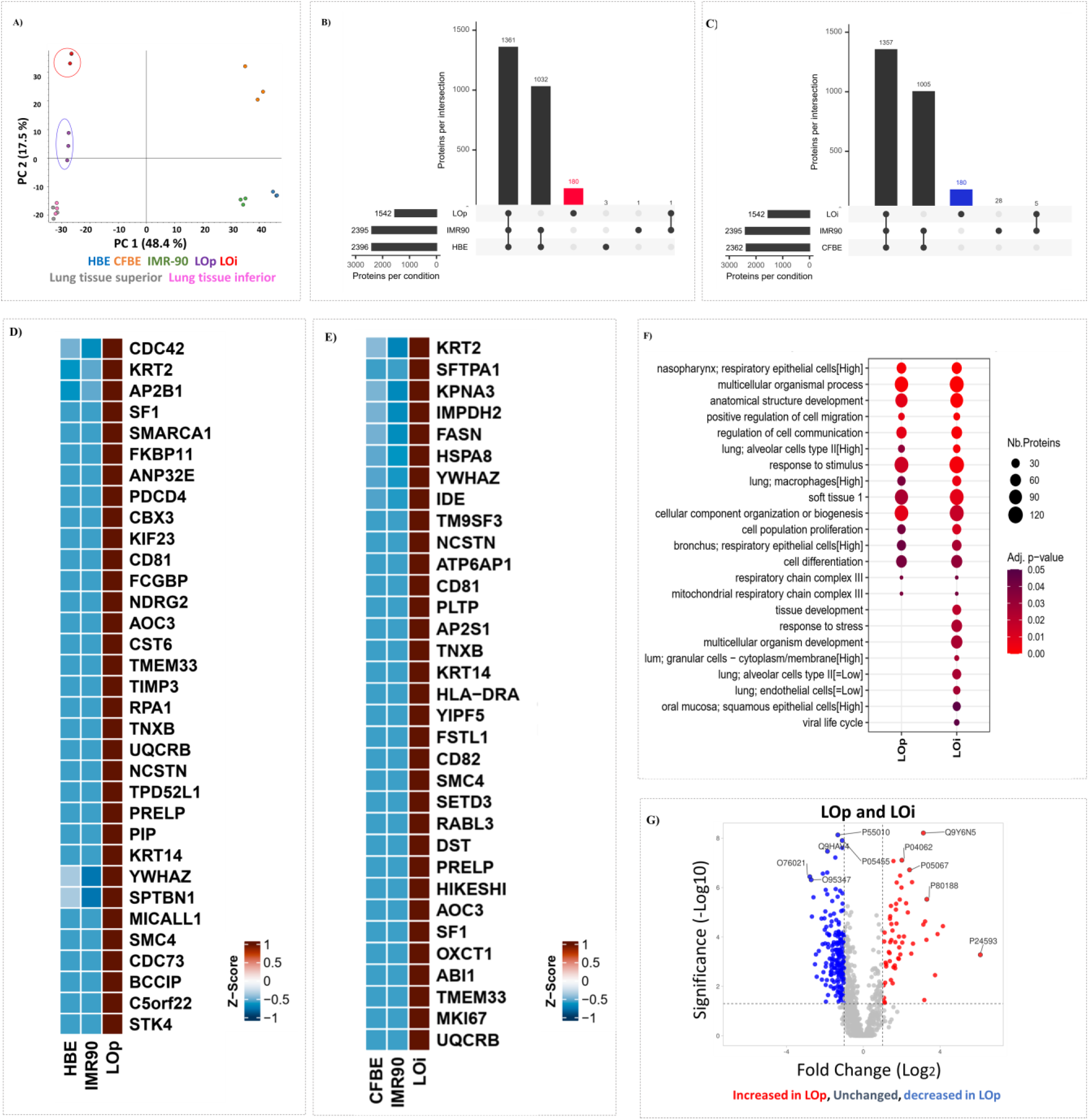
Proteomic profile of lung organoids and cell lines. A) Principal component analysis of cell monolayers, organoids, and tissues. Orange-primary bronchial epithelial cells. Blue-immortalized epithelial cells. Green-Fibroblasts. Purple-Lung organoids derived from primary cells (LOp). Red-Lung organoids derived from immortal cell lines (LOi). Grey-Lung Tissue from superior region. Pink-Lung Tissue from inferior region. B) UpSet plot comparing the number of proteins in LOp (red), IMR-90, and HBE. C) UpSet plot comparing the number of proteins in LOi (blue), IMR-90, and CFBE. D-E) Heatmap showing representative proteins upregulated in LOp, and LOi, respectively, in comparison to cell monolayers from which they are derived from. Grouped by abundances, linked by average. N=3. Upregulation shown in red. Downregulation shown in blue. G) Volcano Plot generated by VolcaNoseR comparing LOp and LOi. Red upregulated proteins in LOp, Grey-unchanged proteins between samples. Blue-downregulated proteins in LOp

Gene ontology was used to confirm the function and the associated tissues of the upregulated proteins in LOi and LOp. Fig. 5F presents a summary of overrepresented terms (set of genes associated with specific functions) categorizing proteins based on their biological roles or properties in LOp and LOi. The figure illustrates the number of proteins corresponding to each term and their statistical significance. Many proteins from both organoid types were associated with lung tissue, soft tissue, respiratory epithelial cells, lung alveolar cells, lung macrophages, and bronchus. For instance, proteins expressed in alveolar cells such as, H3C1, CST6, NDRG2, RPA1, UQCRB, and CXB3 were upregulated in LOp, consistent with the transcriptome findings (Fig.2 and Fig.3), and their distal differentiation.

In organoids derived from immortal cell lines, LOi, proteins associated with viral life cycle, oral mucosa, lung endothelial cells, and response to stress were found (Fig. 5F). In both organoid types, proteins associated with tissue development were found. For instance, terms such as multicellular organismal process, anatomical structure development, biogenesis, and cell migration were all upregulated in the organoids. The data analysis shows that the proteins upregulated in the organoids are not randomly selected; rather, they play key roles in essential functionalities crucial for lung development and function. Together, these findings highlight the targeted and purposeful nature of the protein expression profile observed in the organoids.

Some similarities were found between two types of organoids. The differences between the two types of organoids are shown in Fig. 5G. The proteins upregulated in LOp relative to LOi are shown in red. The proteins upregulated in LOi relative to LOp are shown in blue. Gene ontology and STRING were used to find the functions of the differentiate expressed proteins between the two organoids. The data analysis indicates that LOp has greater specificity to the lung and respiratory system. The proteins upregulated in LOi, in comparison to LOp, were associated with housekeeping gene functions at the DNA level such as DNA repair, DNA metabolic process, and DNA conformational change.

According to their proteomic profile, organoids derived from both cell lines and primary cells exhibit significantly more pertinent proteins than the input cell monolayers from which they are derived. Both organoid types present a higher degree of specificity towards the lung and respiratory system, and they logically display an upregulation of proteins that have diverse and relevant biological functions.

### Proteomic profile of lung organoids vs adult lung tissues

A proteomic analysis of lung organoids (LOp and LOi) and adult lung tissues from three male donors aged 42, 51, and 66. Tissue samples from both superior and inferior regions were included to capture any regional variations in protein expression. The clustering of protein data, as shown in Fig. 6A, resulted in six distinct clusters. These clusters were determined based on the abundance and expression patterns of proteins, employing ward D2 clustering method on Canberra distances. Each cluster represents a group of proteins with similar expression profiles across the samples, suggesting potential functional similarities or shared regulatory mechanisms. Cluster 1 highlights the proteomic similarity between LOp and adult lung tissues. Interestingly, LOp shows resemblance to both lung regions: superior and inferior. Recall, the primary cells are harvested from proximal tissues. The harvest of cells from proximal regions of the tissues (throat and bronchi) can reproduce in part the proteome of further distal tissue, harder and riskier to collect^19,20^. This constitutes a potential benefit for future developments of lung organoids.

**Fig. 6.**
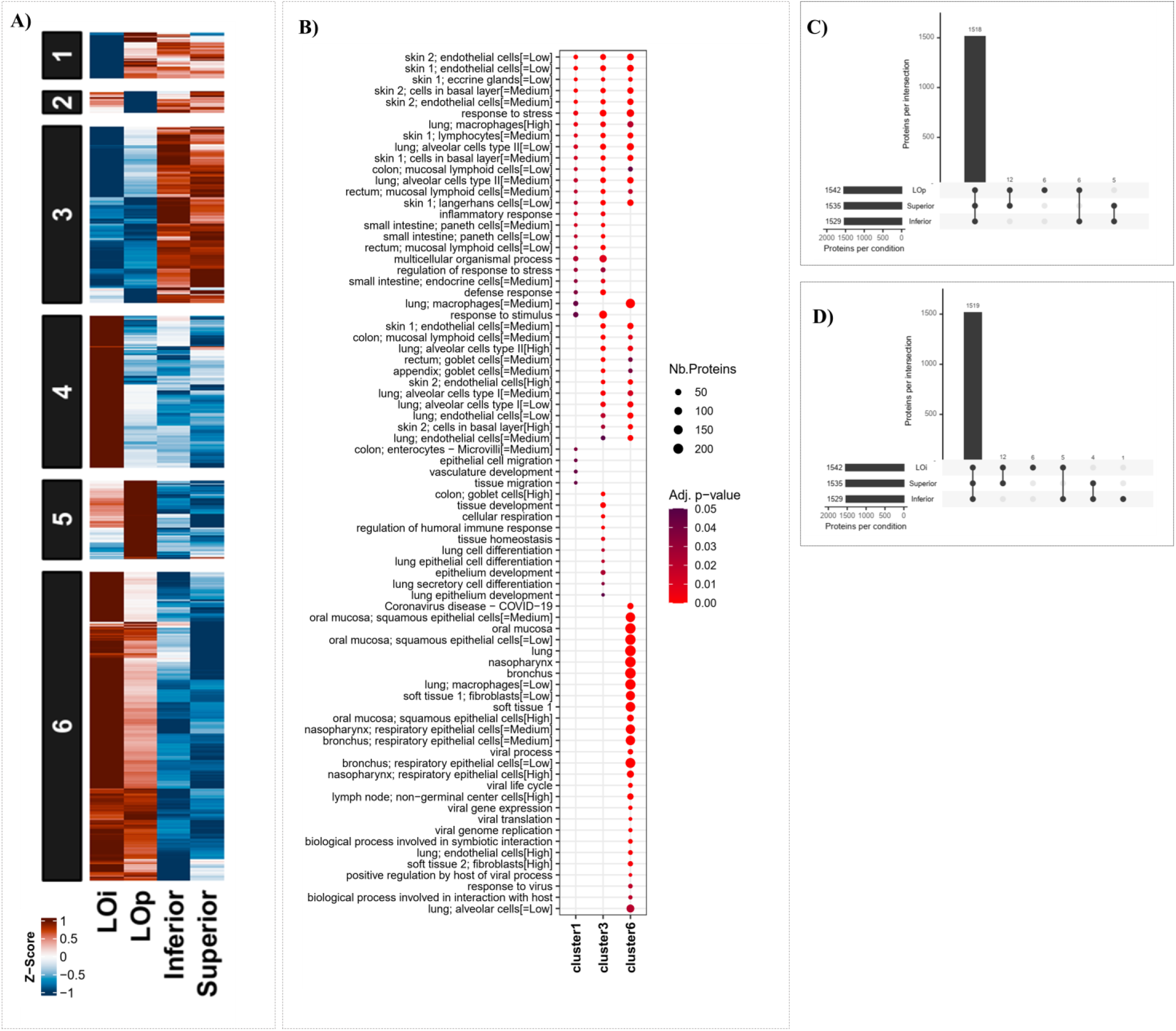
Proteomic profile of lung organoids and adult lung tissues. A) Heatmap of lung organoids derived from immortal cell lines (LOi) and primary cells (LOp) as well as lung tissues from superior and inferior region. Grouped by abundance, linked by average. N=3. Red indicates upregulation of proteins. Blue indicates downregulation. B) Gene Ontology analysis of the overrepresented terms from upregulated proteins in LOp and lung tissues (cluster 1), adult lung tissues (cluster 3) and lung organoids (cluster 6). C-D) UpSet plots visualizing the proteins shared between adult lung tissues and LOp (C) or LOi (D).

Cluster 2, the smallest, illustrates the similarities between the proteome of LOi and adult lung tissues. Cluster 3 is characterized by an upregulation of proteins in adult lung tissues compared to both types of organoids. Conversely, cluster 4 shows a pronounced upregulation of proteins in LOi when compared to LOp and adult lung tissues. Cluster 5 demonstrates an upregulation of proteins in LOp relative to LOi and adult lung tissues. Finally, cluster 6 indicates an overall upregulation of proteins in both organoid types compared to adult lung tissues.

Gene ontology (GO) analysis was conducted on clusters 1, 3, and 6 of the proteome of adult lung tissues and organoids. Figure 6B shows an overview of the terms that are overrepresented in these clusters, according to their biological roles or properties. The data shows the number of proteins associated with each term and its statistical significance. The overrepresented terms in cluster 1 show that the proteins shared between LOp and adult tissues are related to alveolar type II cells, inflammatory response, lung macrophages, basal layer, epithelial migration, microvilli, and response to stress. These results highlight the transregional differentiation of organoids (TDLO-p) into the distal lung region phenotype (AT2 cells). Some phenotypes of the proximal lung region, such as basal cells, are retained. The proteins upregulated in lung tissues with respect to organoids (cluster 3) include overrepresented terms related to goblet cells, indicating a greater number of these cells in the tissue. The functionalities of the proteins upregulated in the tissue include lung epithelial cells, lung cell differentiation, tissue development and secretory cells. On the other hand, the functionalities of the proteins upregulated in the organoids with respect to adult lung tissues (cluster 6) were related to bronchus respiratory epithelium, oral mucosa, and nasopharynx epithelium. This suggests that some of the proteins found more abundantly in the organoids correspond to the proximal lung region. Additionally, several proteins related to viral infectibility were found in the organoids, such as coronavirus disease, virus replication, viral transmission, and response to virus, highlighting the potential application of the model for the study of viral diseases. Numerical analysis was conducted to compare the number of proteins shared between tissues and each type of organoid (Fig.6C-D). As shown in Fig. 6C, adult lung tissue and LOp shared 1518 proteins, from which 5 were exclusive to the tissues, and 6 to the organoids. For the LOi, adult lung tissues and LOi shared 1519 proteins, with only 4 proteins absent in the organoids. LOi organoids had a total of 6 proteins absent from the tissues.

Despite some discrepancies, the proteome of organoids exhibits a notable degree of similarity to that adult lung tissues. Organoids derived from primary cells display proteins from multiple lung regions, underscoring their transregional differentiation potential. Organoids derived from immortal cell lines, show fewer similarities with adult lung tissue. But these LOi offer significant advantages over simple cell lines due to their high specificity to the lung and respiratory system. The LOi bridge the gap between sophisticated organoids and simpler cell lines, providing an alternative to primary cells, which are costlier and more challenging to acquire and culture. Therefore, this offers a more scalable model, named SLO.

### SARS-CoV-2 infection of lung organoids derived from immortalized cell lines (LOi)

One major focus of organoid research is their potential use as presumably accurate model system for infections. The previous sections indicated on the benefits of organoids over cell monolayers, and in terms of protein specificity showed their similarities to adult lung tissues. The organoids generated in this study upregulated proteins associated with viral infections, coronavirus diseases and viral replication (Fig. 6B). Therefore, we aimed to demonstrate possible applications for viral infections, such as SARS-CoV-2, and influenza H1N1. Using RNA sequencing data, genes associated the modulation of critical illness in patients with COVID-19 (SLC6A2, IFNAR1, IFNAR2, OAS3, OAS1, FOXP4, DPP9, TYK2)^53^ were studied in organoids, and adult lung tissues. The genes were expressed in all samples, but interestingly their upregulation was greater in LOi than in LOp, as shown in Fig. 6A. The most abundant gene found in LOi was OAS3. This is an antiviral enzyme which plays a role in cellular innate antiviral response. When activated, it inhibits protein synthesis, thus terminating viral replication^54^.

Interferon induced transmembrane genes were also studied. These genes restrict cellular entry of diverse viral pathogens such as influenza A, COVID-19, and Ebola virus^55^. Interferon Induced Transmembrane 3 (IFITM3) was the most abundant transmembrane in all the samples, as shown in Fig.6B. It was also significantly increased in the LOi.

Entry and priming mechanisms of respiratory viruses such as Angiotensin-Converting-Enzyme 2 (ACE2) and Transmembrane serine protease 2 (TMPRSS2) were also studied. The former is the main mechanism of entry of SARS-CoV-2^56^, while the latter is involved in the priming of SARS-CoV-2^56^, other coronaviruses and influenza^57^. TMPRSS2 was more abundant in LOp than in the tissues and LOi. ACE2 was highly upregulated in tissues, to a lesser degree in LOp, and less abundant in LOi. Furin was also investigated since this protease has the potential to cleave the viral envelope glycoprotein, enhancing viral fusion with host membrane^58,59^. It has been shown in several studies to play an important role in SARS-CoV-2 infectivity^59,60^. Furin was most upregulated in LOi, followed by LOp.

Given that a high number of genes associated with permissibility of respiratory infections were upregulated in the organoids, we hypothesize that these organoids could be used to model SARS-CoV-2 infection. The LOi also upregulated genes associated with the modulation of critical illness of COVID-19 (Fig. 6A), and thus constitute potentially a better model. The presence of the proteins TMPRSS-2 and ACE2 was confirmed in the LOi by immunofluorescence (Fig. 6D-green), and their expression corroborated by qPCR (Fig. 6E). Therefore, we infected three LOi using SARS-CoV-2 with a multiplicity of infection (MOI) of 0.01 and 0.001 for 3 hours as shown in Fig. 6I. The culture medium was then removed in both the upper and lower transwell chambers. Viral aliquot in serum free medium was added in the upper chamber where the organoids were kept for 3 hours at 37°C. They were rocked every 15 minutes. After inoculation, the media was switched to DMEM 2% FBS. At 24 and 48 hours post infection (hpi), the LOi and the cell culture medium were harvested for the detection of viral infection. Our methodology allows the infection of native organoids without their destruction, as for previous studies^24-26^.

No SARS-CoV-2 transcripts were found in the supernatant of negative controls. But SARS-CoV-2 RNA was detected in the supernatants of infected organoids, indicating successful infection. Furthermore, there was significant association between the MOI and the number of viral copies detected, Fig. 6G, indicating that the extent of infection in organoids is sensitive to initial viral load. In addition, organoid lysates were used to evaluate viral replication within the organoid 24 hpi and 48 hpi (Fig.6H). Viral copies were detected only on infected organoids, correlating with the viral copies found in the supernatant. Similarly, organoids show infectibility with influenza H1N1 (S.Fig.6)

## DISCUSSION

### Transregional differentiation

In the present study a novel approach to induce the formation of human lung organoids was invented. The organoids derived from primary cells use cells from trachea/bronchi. These cells are more accessible than deeper lung tissues, the usual source of cells for lung organoids^24-26^. Regardless of their origin, these cells self-assemble into lung organoids, involving upregulation of genes known to be associated with respiratory system development, tube development, and lung alveolus development^25,26^. The expression of multiple cell markers from different regions of the lung shows the transregional differentiation of the model relative to that of organoids representing either proximal or distal lung regions. Overall, the expression of proximal markers of the TDLO-p decreases, apart from basal cells, as the expression of distal markers increases, including surfactant proteins produced by AT2 cells.

The proteome of these organoids demonstrates a similarity to that of adult lung tissues. Furthermore, it supports the transcriptome data by showing the upregulation of several pathways involved in cell differentiation, alveolar development, and respiratory epithelial cells, compared to the input cell lines HBE and IMR-90.

### Scalable lung organoids

The same protocol was successfully adapted to be used with immortal cell lines of bronchial epithelial cells to improve the accessibility of cells and the scalability of the model. These types of cells are cheaper, easier to grow and maintain than their primary counterpart. These organoids, SLO-i, also express several genes and proteins specific to adult lung tissue. Once the organoids are formed cellular markers from proximal regions are upregulated. This highlights the possibility to use the culture microenvironment to recover the cell differentiation capacity of homogenous immortal cells. The microenvironment used was simple, requiring only the 3D culture of fibroblasts within a hydrogel that mimics natural stiffness, along with ECM proteins. Neither an ALI culture, tidal breathing, nor vascularization were used. Enhancing the culture microenvironment in the future could lead to higher cell differentiation into distal lung regions. The differentiation of LOi into markers associated with distal lung regions such as HOPX for AT1 were not significatively upregulated, which suggest less heterogeneity and less cell transregional differentiation of the model in comparison to TDLO-p.

The proteome of SLO-i included large pool of proteins specific to the respiratory system, including lung, and trachea, as well as immune system proteins, which may be beneficial to model viral infections. The data establishes the superiority of organoids over cell monolayers in terms of specificity, while requiring simple culture methods during organoid formation.

### Tubular morphology of lung organoids

The new lung organoid models, TDLO-p and SLO-i, exhibit a more sophisticated branching morphology compared to spheroidal lung organoids. Typically, lung organoids derived from iPSCs have a tubular morphology^17,22^. This study is the first to achieve such morphology using adult stem cells, in the case of TDLO-p, and immortalized cells, in the case of SLO-i. Typically, there are at least two levels of branching observed in the organoids, with some variations depending on the donor. For instance, organoids derived from primary cells obtained from female donors tend to exhibit more branching structures compared to those from male donors. This achieved morphology is a simplified version of the lower airways, since this one form a branching tree of 23 generations^61^.

Based on enrichment functional analysis, the morphology of the organoids derived form primary cells (TDLO-p) appears to be the result of well orchestrated processes, as evidenced by the upregulation of genes associated with tube development, and branching structures. We think that this morphology is the result of using lung fibroblasts as feeder cells, since their removal from the protocol yields a spheroidal morphology in SLO-i.

This constitutes an important step in the self-organization of cells into a functional organ. Microscopic examinations of SLO-i, reveal mimetic self-organization traits, such as pseudostratified squamous epithelium^62^, and microvilli structures^47^, but also cell debris deposited randomly within void structures.

### Genetic markers and proteins associated with viral infectability in organoids

Scalable lung organoids show upregulation of proteins associated with viral life cycle in respect to the input cell lines, CFBE and IMR-90, used to form the SLO-i. These viral life cycle proteins include those involved in viral entry, latency, transcription, and replication of the virus, as well as proteins associated with host virus interactions^63^. This was confirmed by the upregulation of ACE2 and TMPRSS2 in the organoids, compared to the expression on CFBE grown in ALI. Both ACE2 and TMPRSS2 are essential for SARS-CoV-2 infection^56^.

Both organoids, TDLO-p and SLO-i show upregulation of proteins associated with viral infectability in respect to adult lung tissues. These proteins include specific terms associated with COVID-19 disease, viral replication, viral process, viral life cycle, viral translation, viral genome replication, response to virus, and positive regulation by host-viral process (Fig. 6B). These data align with the results obtained from transcriptomics, which showed upregulation of specific genes involved in pathways necessary for COVID-19 and influenza infection (Fig. 7A-C).

**Fig. 7.**
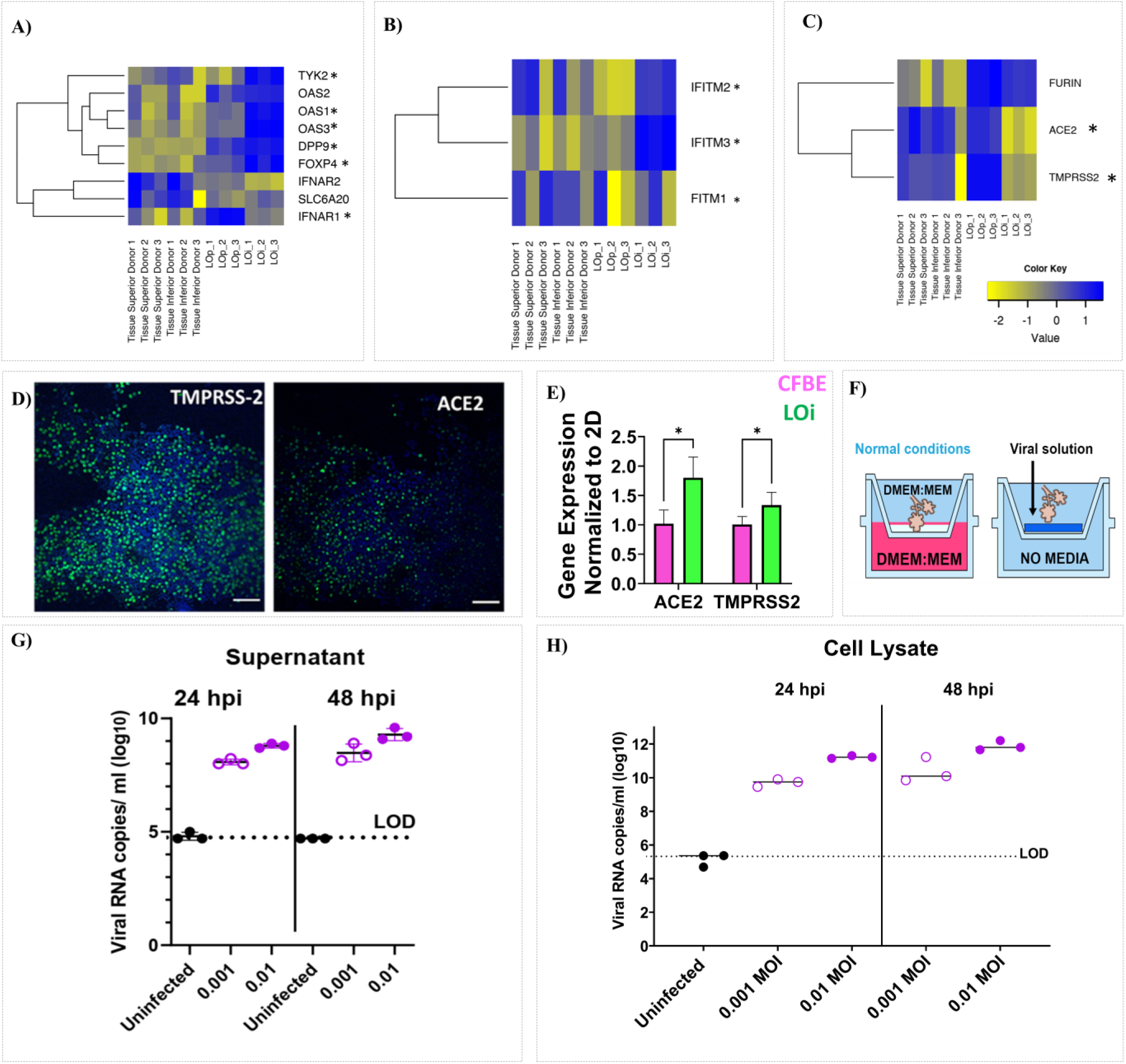
Exploring SARS-CoV-2 infection on lung organoids. A-C) RNA sequence data showing heatmap of genes associated with A) critical illness of COVID-19, B) Interferon induced transmembrane. C) Entry receptors, priming and cleavage proteins for SARS-CoV-2. T-tests (LOp and LOi) * p. adjust<0.05. N=3. Upregulation shown in blue. D) Images of LOi organoids at day 7 expressing in green TMPRSS-2 and ACE-2, blue-nuclei. Magnification 10X. Scale Bar 100 μm. E) Gene expression by qPCR of TMPRSS-2 and ACE2 in organoid-green and CFBE monolayers-pink. n=3, p<0.05. F) Schematic representation of viral infection in the organoids. G) Viral copies /ml (log10) on supernatant of organoids under MOI: 0.01 and 0.001 SARS-CoV-2 infected-purple, and non-infected-black, 24 hpi and 48 hpi (n=3). H) Viral copies/ml (log10) on cell lysate of organoids MOI: 0.01 and 0.001 SARS-CoV-2 infected-purple, and non-infected-black, 24 hpi and 48 hpi (n=3). Two-way ANOVA p<0.05 in every condition. LOD: Limit of detection.

These findings suggest that TDLO-p and SLO-i are relevant models to study respiratory viral infections. This will not only allow for a greater understanding of infectious respiratory disease etiology, but also pave the way for using these organoids as a tool to discover new and effective anti-viral therapies.

### Viral infection in scalable lung organoids

Given the low cost and availability of organoids derived from immortal cells, SLO-i was infected with SARS-CoV-2 and influenza H1N1. The model responded to the infection, as indicated by the detection of viral RNA copies in its supernatant and cell lysate. Unlike other models, the organoids did not need to be destroyed to become infected^14,16,21^. This is a remarkable aspect as it allows the study of viral infections in a 3D environment, preventing possible viral mutations^3^.

Ongoing studies are directed at the infection of TDLO-p with influenza H1N1 and SARS-CoV-2, alongside the functional analysis of SLO-i using pre-approved drugs. Functional enrichment analysis after infection is being conducted to compare the expression of genes with data from clinical databases of patients with SARS-CoV-2.

Current research efforts also include comparing the phenotypes of TDLO-p and SLO-i with other previously reported adult lung organoids representing either proximal or distal lung^15^ region or a more heterogeneous lung model recently reported^16^.

## LIMITATIONS OF THE STUDY

The new lung organoids have a tubular morphology with some branching structures. They appear to be more similar to adult lungs than spheroidal lung models. They remain a much-simplified representation of the complex morphology of adult lungs^61^. Markers of proximal and distal regions of the lung remark the transregional differentiation of LOp, and its proteomic profile shows its similarity that of adult lung tissues. But the spatial relationship of proximal and distal markers is so far randomly distributed, which may be a consequence of the size of the organoid in comparison to the actual organ. Future work is needed to gain insights into the structure of the organoid using more qualitative methods than the ones here provided. Single-cell sequencing can provide in-depth characterization to assess cellular composition and track their changes upon modification of the protocol.

The proteome of the organoids included several proteins associated with the response to mechanical stresses. Therefore, the model would benefit from the integration of mechanical forces which occur during breathing. Proteins from the innate immune system were upregulated during the organoid formation, which may support the physiological relevance of the model for viral infections. But the incorporation of innate immune cells, such as macrophages would be beneficial to better understand the pathophysiology of respiratory infections, and cell-cell communication.

## ACKNOWLEDGMENTS

We thank Dr. Guangyu Bao for his guidance and training in rheology. Genome Quebec performed RNA sequencing. We thank Virginie Calderon and the Montreal Clinical Research Institute (IRCM) for the bioinformatics analysis of RNA sequencing data. We thank the Proteomics and molecular Analysis Platform at Research Institute of the McGill University Health Centre (RI-MUHC) for performing tandem mass tags and for the statistical analysis. Specifically, we thank Jennifer Nedow, Lorne Taylor and Jenna Cleyle. We thank Charlotte Fouquet for her help extracting RNA from some samples infected with SARS-COV-2. This work was supported by NIH fund RO1 DC018577 (Mongeau, PI). The views expressed in the present publication are the authors’ own and do not necessarily represent those of the NIH. We thank CONACYT for supporting the doctoral studies of Alicia Reyes Valenzuela.

## CONTRIBUTIONS

Alicia Reyes Valenzuela conceived and designed the project, performed the experiments, developed and optimized the protocol for organoid formation, fabricated and prepared the organoids, and analyzed the data (unless stated otherwise). Mark Turner along with Prof. John Harahan designed primers, contributed to qPCR analysis, and contributed their expertise in the culture of epithelial primary cells. Prof. John Hanrahan provided the ethical protocol for the use of human cell lines. Christophe Lachance-Brais edited and reviewed the manuscript. Prof. Hojatollah Vali obtained images using Focused Ion Beam microscopy. Nathan Markarian performed the organoid infection with SARS-CoV-2, performed RNA extraction of infected organoids, and analyzed viral RNA copies in the organoid lysate and supernatant. Prof. Silvia Vidal designed the experiments for viral infection and provided guidance for the viral study. Prof. Luc Mongeau supervised the study, reviewed the manuscript, and provided guidance and oversight.

## STAR METHODS

### KEY RESOURCES TABLES

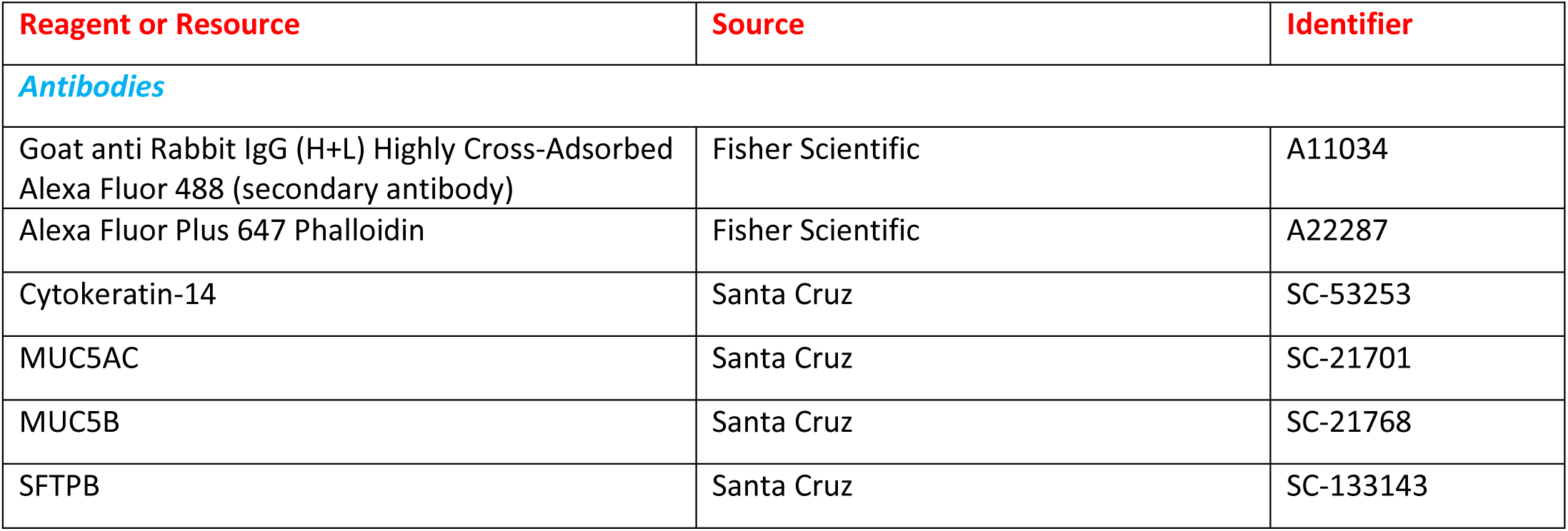
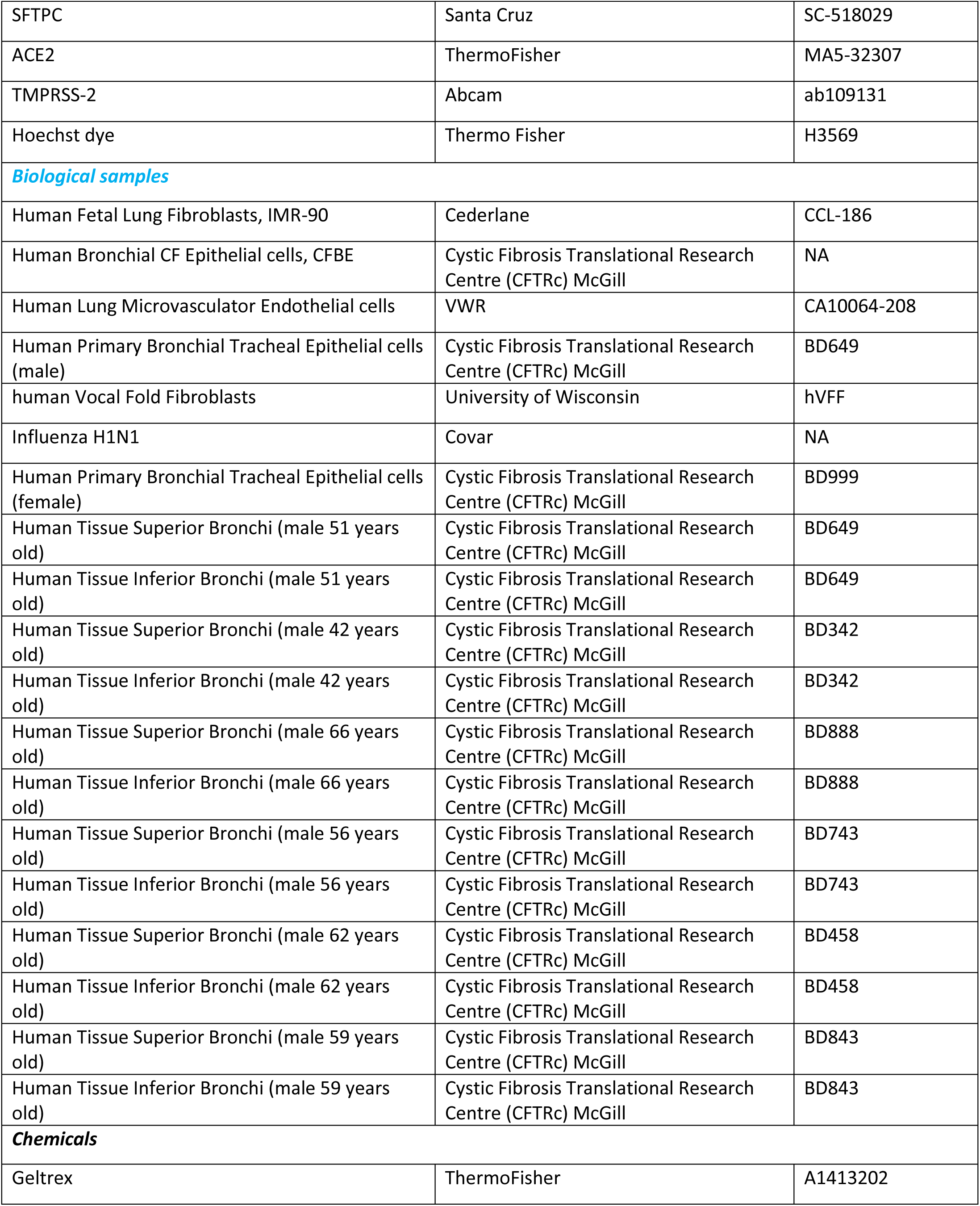
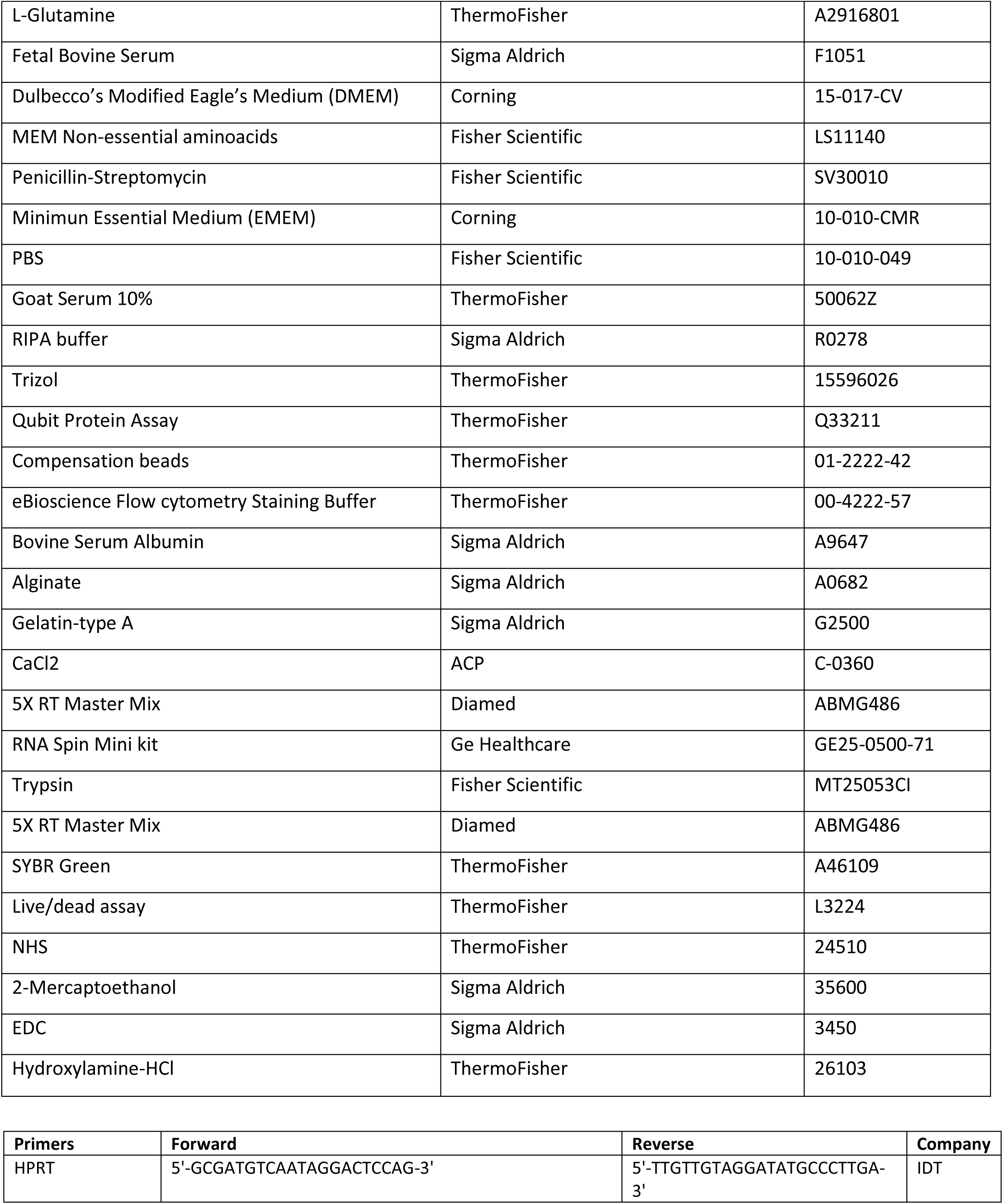
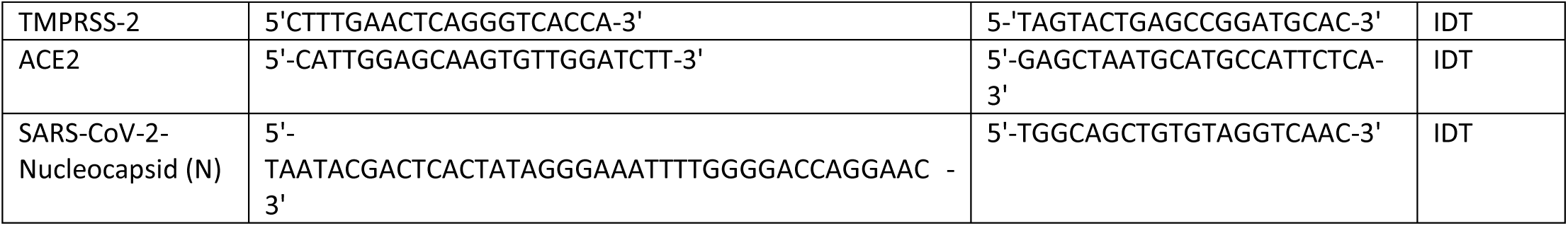

### METHODS DETAILS

#### Cell culture and CFBE in air-liquid interface

CFBE and IMR-90 were cultured in Eagle’s minimum essential medium (VWR, CA45000-380) supplemented with 10% (vol/vol) FBS (Sigma-Aldrich, F1051), 2 mM L-glutamine (Thermo Fisher, A2916801), 1% Penicillin Streptomycin (Fisher Scientific, SV30010), and incubated at 37°C in 5% CO_2_-95% O_2_. Human Vocal Fold Fibroblasts (hVFF) were cultured in Dulbecco’s Modified Eagle’s Medium (Life Technologies, 12430054) supplemented with 10% (vol/vol) FBS (Sigma-Aldrich, F1051), 1% Aminoacids solution (Thermo Fisher, 11130051), 1% Penicillin Streptomycin (Fisher Scientific, SV30010), and incubated at 37°C in 5% CO_2_-95% O_2_. For establishment of ALI, CFBE were seeded on fibronectin (Promo cell, C-43060) coated permeable cell culture inserts (Fisher Scientific, 08-771-10) and cultured for 21 days before the study.

#### Primary airway epithelium culture in air-liquid interface

Primary human bronchial epithelial were provided by Primary Airway Cell Biobank (PACB) in the Cystic Fibrosis Translational Research Centre at McGill University. Cell from male and female donors of 51 and 53 years old, respectively were used. Cells were seeded on collagen IV coated cell culture inserts (Fisher Scientific, 08-771-10) and cultured in ALI medium provided by PACB. After 3 days, the apical medium is removed, and cells were allowed to differentiate for 28 days at the air-liquid interface before study. The medium in the basolateral chamber is changed every two days. Once a week, the apical mucus is aspirated carefully to avoid disruption of the monolayer. Followed by washing the apical side with PBS without Ca^2+^.

#### Establishing adult CFBE-derived human lung organoids

Human fetal lung, IMR90, cells were cultured overnight in different alginate-gelatin hydrogels in a density of 250,000 cells per/cm^2^ using DMEM (Corning, 15-017-CV) with Penicillin-Streptomycin (Fisher Scientific, SV30010) non-essential aminoacids (Fisher Scientific, LS11140050) and FBS (Sigma-Aldrich, F1051) at 37°C. The media was removed, and the hydrogels were coated using 100 μl/cm^2^ of Geltrex (Thermo Fisher, A1413202) during 30 minutes at 37°C. Afterwards Human Bronchial CF Epithelial cell line was seeded on the top in a density of 500,000 cells per/cm^2^ using EMEM (Corning) with L-Glutamine (Thermo Fisher, A2916801) forming a co-culture interface. The next day the medium was substituted by a combination in a ration 1:1 of DMEM and EMEM supplemented with 3 mM CaCl_2_ (Sigma, Aldrich, C4901). The organoids were formed by cellular and extracellular matrix interactions. After 5 days of culture, the maturation, and branching of the organoids was observed, and the organoids were cultured in submerged until day 21 at 37°C under a humidified atmosphere with 5% CO_2_ and 95% air. During culturing, medium was refreshed at most every two days.

#### Immunofluorescence Staining

##### Morphology

The morphological analysis of the organoids was made at day 7, 16, 21, 30 and 40 days, according to the study. Organoids were washed with PBS (Fisher Scientific, 10-010-049) supplemented with 10 mM CaCl_2_ (Sigma, Aldrich, C4901) three times for 3 minutes. Afterwards, the organoids were fixed with 3.7% paraformaldehyde, permeabilized with 0.3% Triton X-100 (Sigma Aldrich, T9284), and blocked for 2 hours with 10% Goat serum (Thermo Fisher, 50062Z) at room temperature. Organoids were culture with primary antibodies diluted in 10% goat serum at 4°C overnight followed by incubation of secondary antibody Goat-anti Rabbit IgG Alexa Fluor 488 (Fisher Scientific, A11034), for 2 hours at room temperature. Nuclei and actin filaments were counterstained with Hoechst dye (Thermo Fisher H3569) and Phalloidin-647 (Fisher Scientific, A22287) respectively. The confocal images were acquired using an LSM-710 inverted confocal microscope.

##### Protein expression

To evaluate the expression of proteins related to infection and replication of SARS-COV-2, such as Transmembrane Protease Serine 2, TMPRSS, and Angiotensin Converting Enzyme 2, ACE2, primary antibodies were used and incubated overnight in 10% goat serum followed by incubation of secondary antibody Goat-anti Rabbit IgG Alexa Fluor 488 (Fisher Scientific, A11034) for 2 hours at room temperature. The primary antibodies used were the ACE2 recombinant rabbit monoclonal antibody (SN0754) from ThermoFisher, and Recombinant Anti-TMPRSS-2 antibody (AB109131) from Abcam.

#### Focused Ion Beam microscopy

Organoids derived from immortalized cells at 21 days of culture were embedded in resin after sequential fixation in 2.5% glutaraldehyde in 0.1 M cacodylate buffer (pH 7.2) at 4 °C. The sections were examined under 2.5 nm at 30 kV Focused Ion Beam Microscope.

#### RNA isolation and quantitative PCR

RNA was extracted using Illustra RNA Spin Mini kit (Ge Healthcare, GE25-0500-71) according to the manufacturer’s instructions. RNA (500 ng) was reverse transcribed with 5X RT Master Mix (Diamed, ABMG486) with incubation at 25°C for 10 minutes, 42°C for 1 h, and 85°C for 5 minutes. Quantitative PCR (qPCR) was performed using Power track SYBR green (ThermoFisher, A46109). The reaction was performed using a QuantStudio 7 Flex Real-Time PCR system and the following protocol: 20 s at 95°C and 40 cycles at 95°C (1 s) and 60°C (20 s). The change in gene expression relative to housekeeping gene (ΔΔCt) was calculated using the manufacturer’s software package. Primers were predesigned for IDT, except for the house keeping gene HPRT, ACE2, and TMPRSS2. They are listed in key Resources Table II.

#### Bulk RNA sequencing and analysis

The extraction, library preparation and sequencing services were provided by the Génome Québec Centre d’expertise et de services. RNA extraction was performed according to the sample type, as follows.

*<u>Lung</u>*, RNeasy Plus Universal mini kit (QIAGEN) was used. Frozen tissue samples were weighed and processed on ice to prevent thawing. For homogenisation: 900 μl of RNeasy Plus Universal Mini Kit provided lysis buffer reagent (i.e. QIAzol) was added to the tissues and homogenized using a QIAGEN TissueLyser II with 2,8 mm ceramic beads, for 2 cycles of 30 Hz x 2 minutes plus 1 cycle of 30 Hz x 1 minute. Extraction was performed according to the manufacturer’s instructions. RNA was eluted in 35 μl buffer provided with the extraction kit. RNA quality was determined by the RNA Integrity Number (RIN), measured by 2100 Bioanalyzer (Agilent Technologies) using RNA 6000 Nano kit, following the manufacturer’s protocol.

*<u>Organoid,</u>* three organoids were disassociated an integrated as one sample to obtain enough RNA per sample. This was done on three independent sets of organoids. Therefore 9 organoids were processed and divided in 3 samples. RNeasy Plus Universal mini kit (QIAGEN) was used. For homogenisation 900 μl Trizol reagent was added to the samples and homogenized using a QIAGEN TissueLyser II with 2,8 mm ceramic beads, for 3 cycles of 30Hz x 2 minutes. Extraction was performed according to the manufacturer’s instructions. RNA was eluted in 35 μl buffer provided with the extraction kit.

RNA quality was determined by the RNA Integrity Number (RIN), measured by 2100 Bioanalyzer (Agilent Technologies) using RNA 6000 Nano kit, following the manufacturer’s protocol.

*<u>Cells,</u>* miRNeasy mini kit (QIAGEN) was used. Cell samples were homogenized in the TRIzol lysis reagent by passing the lysate 10 times through a 20-gauge needle attached to a sterile 3 ml plastic syringe.

Extraction was performed according to the manufacturer’s instructions. RNA was eluted in 35 μl buffer provided with the extraction kit. RNA quality was determined by the RNA Integrity Number (RIN), measured by 2100 Bioanalyzer (Agilent Technologies) using RNA 6000 Nano kit, following the manufacturer’s protocol.

##### Library preparation

Ribosomal RNA was depleted from 250 ng of total RNA using QIAseq FastSelect. cDNA synthesis was achieved with the NEBNext RNA First Strand Synthesis and NEBNext Ultra Directional RNA Second Strand Synthesis Modules (New England BioLabs). The remaining steps of library preparation were done using and the NEBNext Ultra II DNA Library Prep Kit for Illumina (New England BioLabs). Adapters and PCR primers were purchased from New England BioLabs.

##### Library QC

Libraries were quantified using the KAPA Library Quanitification Kits - Complete kit (Universal) (Kapa Biosystems). Average size fragment was determined using a LabChip GX II (PerkinElmer) instrument.

##### Sequencing

The libraries were normalized and pooled and then denatured in 0.02N NaOH and neutralized using HT1 buffer. The pool was loaded at 175 pM on a Illumina NovaSeq S4 lane using Xp protocol as per the manufacturer’s recommendations. The run was performed for 2x100 cycles (paired-end mode). A phiX library was used as a control and mixed with libraries at 1% level. Base calling was performed with RTA v3. Program bcl2fastq2 v2.20 was then used to demultiplex samples and generate fastq reads.

##### Analysis

The quality of the raw reads was assessed with FASTQC v0.11.8. After examining the quality of the raw reads, no trimming was deemed necessary. The reads were aligned to the GRCh38 human reference genome with STAR v2.7.6a, with an average of 90 % of reads uniquely mapped. The raw counts were calculated with FeatureCounts v1.6.0 based on the human reference genome (release 108). Differential expression was performed using DESeq2 R package. Differentially expressed genes were define using adj-pvalue < X and |log2FCshrink|≥4 X leading respectively X, Y, Z DEGs for A vs B, A vs C and B vs C. DEGs heatmap were drawn based on z-score of normalized count. Bioinformatics analyses were performed at the Bioinformatics core facility from Montreal Clinical Research Institute (IRCM).

#### Tandem Mass Tag

##### Sample preparation

Cells monolayers and organoids were washed with cold PBS (X3) following by incubation on ice cold RIPA buffer (Sigma, R0278). The cell monolayers and organoids were broken using a plastic cell scraper. The organoids were stabbed using a 21G needle until completely disassociated. Cells were collected and centrifuged at 16000 G for 50 minutes at 4°C. The supernatants were collected, protein quantification was performed with Qubit Protein Assay (ThermoFisher, Q33211), and normalized to 45 ng of protein. Samples were stored at -80 °C until analysis. All the reported proteins have a 95% threshold.

##### Experimental procedure

Samples were treated with TMT-16plex reagents (ThermoFisher Scientific) according to the manufacturer’s instructions. Labelled peptides were fractionated using Pierce™ High pH Reversed-Phase Peptide Fractionation Kit into 8 fractions.

Each fraction was re-solubilized in 0.1% aqueous formic acid and 2 micrograms of each was loaded onto a Thermo Acclaim Pepmap (Thermo, 75uM ID X 2cm C18 3uM beads) precolumn and then onto an Acclaim Pepmap Easyspray (Thermo, 75uM X 15cm with 2uM C18 beads) analytical column separation using a Dionex Ultimate 3000 uHPLC at 250 nl/min with a gradient of 2-35% organic (0.1% formic acid in acetonitrile) over three hours running the default settings for MS3-level SPS TMT quantitation (McAlister et al, 2014 - Anal Chem. 2014 Jul 15;86(14):7150-8. doi: 10.1021/ac502040v.), on an Orbitrap Fusion instrument (ThermoFisher Scientific) was operated in DDA-MS3 mode.

Briefly, MS1 scans were collected at 120,000 resolutions, scanning from 375-1500 m/z, collecting ions for 50 ms or until the AGC target of 4e5 was reached. Precursors with a charge state of 2-5 were included for MS2 analysis, which were isolated with an isolation window of 0.7 m/z. Ions were collected for up to 50 ms or until an AGC target value of 1e4 was reached and fragmented using CID at 35% energy; these were then read out on the linear ion trap in rapid mode. Subsequently, the top 10 (height) sequential precursor notches were selected from MS2 spectra for MSquantitative TMT reporter ion analysis, isolated with an m/z window of 2 m/z and fragmented with HCD at 65% energy. Resulting fragments were read out in the Orbitrap at 60,000 resolutions, with a maximum injection time of 105 ms or until the AGC target value of 1e5 was reached.

##### Mass Spectrometry Raw Data Analysis

To translate .raw files into protein identifications and TMT reporter ion intensities, Proteome Discoverer 2.2 (ThermoFisher Scientific) was used with the built-in TMT Reporter ion quantification workflows. Default settings were applied, with Trypsin as enzyme specificity. Spectra were matched against the humn protein fasta database obtained from Uniprot (2022). Dynamic modifications were set as Oxidation (M), and Acetylation on protein N-termini. Cysteine carbamidomethyl was set as a static modification, together with the TMT tag on both peptide N-termini and K residues. All results were filtered to a 1% FDR.

##### Analysis

The set of proteins for each group was analyzed using the STRING software database available online (https://string-db.org/). STRING collects and scores evidence from several sources, including text mining from the scientific literature, databases of interaction experiments and complex pathways, computational interactions, and predictions from co-expression ^30^. Only nodes and networks with high strength (p<0.05) and low false discovery rate are shown.

Protein abundances across different conditions were collected in a matrix, and unidentified proteins were imputed with a value of zero. A pseudo-count of one was then added to the matrix, and a z-score was calculated for each protein (row-wise). Heatmaps were created in R with the ComplexHeatmap package, employing the ward. D2 clustering method on Canberra distances. Clusters of proteins were extracted, and overrepresentation analyses were performed on Gene Ontology (GO) terms, KEGG pathways, Human Protein Atlas (HPA) tissues, and CORUM complexes with the R package gprofiler2 (PMID 31066453). The p-values were statistically corrected with the false discovery rate (FDR) correction method. Statistically significant terms with adjusted p-values < 0.05 were selected and presented in dotplots using the R package ggplot2. Additionally, UpSet plots were created in R with the upsetR package to enumerate non-redundant interactions identified among the different condition

#### Rheological measurements

The shear moduli of AlGe hydrogels were determined using a torsional rheometer with parallel plates (Discovery HR-2). Isothermal time sweeps were applied at a frequency of 0.1 Hz and 0.1 % strain at 37 °C. The shear moduli were recorded and analyzed using TA instruments software. Experiments were conducted in triplicate.

#### Biomaterial synthesis

*AlGe-*A solution of 2% Alginic acid molecular weight (MW) 212.121 g/mol (Sigma, A0682) was prepared in MilliQ water and sterilized with 0.2 μm PES filter (Fisher Scientific, FB12566502). The solution was mixed with 5% sterile Gelatin type A (Sigma, G2500). The hydrogel was formed by incubation with 100 mM CaCl_2,_ MW 147.02 g/mol (ACP, C-0360) for 15 minutes.

#### SARS-CoV-2 infection in human lung organoids

A clinical viral isolate (isolated prior to November 2020) termed SARS-CoV-2 CP13.32 passage 2 (Genbank accession no. 599736; lineage B1.1.147) stock was obtained from the McGill University Health Center and it was propagated in VeroE6 cells five times. The latter stock (P5) was used for infecting the organoid samples.

Before infection, the organoids were washed twice with plain DMEM. Organoids were then inoculated with MOI of 0.001 and 0.01 of SARS-CoV-2 (CP13.32 strain, P5) in a serum-free medium for 3 hours at 37°C and were rocked every 15 minutes. After inoculation, the media was switched to DMEM 2% FBS. At the indicated times, the organoids and cell culture medium were harvested for detection of viral load. The cell-free culture medium was analyzed as “supernatant.”

#### Quantification of SARS-CoV-2 RNA in Supernatant using RT–qPCR

At the indicated times of infections, viral RNA was extracted from 140 μL of supernatant using the QIAamp Viral Mini Kit. The primers used are described in the supplementary section. SARS-CoV-2 RNA was detected in a 10µL reaction containing 1 µL of extracted RNA sample, 200nM of SARS-CoV-2-specific primers, 1X RT enzyme mix in the reaction mixture containing SYBR Green (*Power* SYBR™ Green RNA-to-C_T_™ *1-Step* Kit). The one-step RT–qPCR comprises a 30min RT step at 38 °C and 10 min of AmpliTaq Gold DNA polymerase activation at 95 °C, followed by 40 cycles of PCR at 95 °C for 15 s and 60 °C for 1 min. The RNA copy number was calculated from a standard curve 10-fold serially diluted RNA made from in vitro transcripts consisting of SARS-CoV-2 with known copy numbers. The limit of detection (LOD) was 51 genomic copies per reaction.

#### Quantification of SARS-CoV-2 RNA in Cell Lysates using RT–qPCR

At the indicated times of infections, viral RNA was extracted from cell lysates (organoids resuspended in 1mL of TRIzol) using the PureLink RNA Mini Kit (ThermoFisher). The primers used are described in the supplementary section. SARS-CoV-2 RNA was detected in a 10µL reaction containing 1 µL of extracted RNA sample, 200nM of SARS-CoV-2-specific primers, 1X RT enzyme mix in the reaction mixture containing SYBR Green (*Power* SYBR™ Green RNA-to-C_T_™ *1-Step* Kit). The one-step RT–qPCR comprises a 30min RT step at 38 °C and 10 min of AmpliTaq Gold DNA polymerase activation at 95 °C, followed by 40 cycles of PCR at 95 °C for 15 s and 60 °C for 1 min. The RNA copy number was calculated from a standard curve 10-fold serially diluted RNA made from in vitro transcripts consisting of SARS-CoV-2 with known copy numbers. The limit of detection (LOD) was 48.6 genomic copies per reaction.

#### Statistical analysis

For transcriptomic data, differential expression analysis was performed using the DESeq2 R package. Differentially expressed genes (DEGs) were defined using an adjusted p-value < X and |log2FCshrink| ≥ 4, resulting in X, Y, and Z DEGs for A vs B, A vs C, and B vs C, respectively. For specific genes (e.g., six major cell types, critical illness of COVID-19), a t-test was performed between groups, with significance considered at p.adj < 0.05.

For proteomic data, the p-values were statistically corrected with the false discovery rate (FDR) correction method. Statistically significant terms with adjusted p-values < 0.05 were selected and presented in dotplots using the R package ggplot2.

For the rest of the data, GraphPad was used for statistical analysis. According to the type of data, either two-way ANOVA or t-test was performed. P<0.05 was consider statistically significant. *p<0.05, **p<0.01, ***p<0.005

## REFERENCES

1. Chen, D. et al. Human Organoids as a Promising Platform for Fighting COVID-19. Int. J. Biol. Sci. 18, 901–910 (2022).

2. Du, J. et al. Stem cell therapy: a potential approach for treatment of influenza virus and coronavirus-induced acute lung injury. Stem Cell Res. Ther. 11, 192 (2020).

3. Porotto, M. et al. Authentic Modeling of Human Respiratory Virus Infection in Human Pluripotent Stem Cell-Derived Lung Organoids. mBio 10, e00723–19, /mbio/10/3/mBio.00723-19.atom (2019).

4. World Health Organization. Tuberculosis. https://www.who.int/news-room/fact-sheets/detail/tuberculosis (2023).

5. Si, L. et al. Human Organ Chip-Enabled Pipeline to Rapidly Repurpose Therapeutics during Viral Pandemics. http://biorxiv.org/lookup/doi/10.1101/2020.04.13.039917 (2020).

6. Kim, J., Koo, B.-K. & Knoblich, J. A. Human organoids: model systems for human biology and medicine. Nat. Rev. Mol. Cell Biol. 21, 571–584 (2020).

7. Hou, Y. J. et al. SARS-CoV-2 Reverse Genetics Reveals a Variable Infection Gradient in the Respiratory Tract. Cell 182, 429–446.e14 (2020).

8. Kalil, A. C. & Thomas, P. G. Influenza virus-related critical illness: pathophysiology and epidemiology. Crit. Care 23, 258 (2019).

9. Ramezankhani, R., Solhi, R., Chai, Y. C., Vosough, M. & Verfaillie, C. Organoid and microfluidics-based platforms for drug screening in COVID-19. Drug Discov. Today 27, 1062–1076 (2022).

10. Mulay, A. et al. SARS-CoV-2 infection of primary human lung epithelium for COVID-19 modeling and drug discovery. Cell Rep. 35, 109055 (2021).

11. Sprott, R. F. et al. Flagellin shifts 3D bronchospheres towards mucus hyperproduction. Respir. Res. 21, 222 (2020).

12. Cunniff, B., Druso, J. E. & van der Velden, J. L. Lung organoids: advances in generation and 3D-visualization. Histochem. Cell Biol. 155, 301–308 (2021).

13. Rock, J. R. et al. Basal cells as stem cells of the mouse trachea and human airway epithelium. Proc. Natl. Acad. Sci. 106, 12771 (2009).

14. Zhou, J. et al. Differentiated human airway organoids to assess infectivity of emerging influenza virus. Proc. Natl. Acad. Sci. 115, 6822–6827 (2018).

15. Chiu, M. C. et al. A bipotential organoid model of respiratory epithelium recapitulates high infectivity of SARS-CoV-2 Omicron variant. Cell Discov. 8, 57 (2022).

16. Tindle, C. et al. Adult stem cell-derived complete lung organoid models emulate lung disease in COVID-19. eLife 10, e66417 (2021).

17. Miller, A. J. et al. Generation of lung organoids from human pluripotent stem cells in vitro. Nat. Protoc. 14, 518–540 (2019).

18. Konishi, S. et al. Directed Induction of Functional Multi-ciliated Cells in Proximal Airway Epithelial Spheroids from Human Pluripotent Stem Cells. Stem Cell Rep. 6, 18–25 (2016).

19. Chiu, Y.-W. et al. Costs of Biopsy and Complications in Patients with Lung Cancer. Clin. Outcomes Res. Volume 13, 191–200 (2021).

20. Hutchinson, J. P., Fogarty, A. W., McKeever, T. M. & Hubbard, R. B. In-Hospital Mortality after Surgical Lung Biopsy for Interstitial Lung Disease in the United States. 2000 to 2011. Am. J. Respir. Crit. Care Med. 193, 1161–1167 (2016).

21. Sachs, N. et al. Long-term expanding human airway organoids for disease modeling. EMBO J. 38, e100300 (2019).

22. Chen, Y.-W. et al. A three-dimensional model of human lung development and disease from pluripotent stem cells. Nat. Cell Biol. 19, 542–549 (2017).

23. Dye, B. R. et al. Human lung organoids develop into adult airway-like structures directed by physico-chemical biomaterial properties. Biomaterials 234, 119757 (2020).

24. Sang, Y., Miller, L. C., Nelli, R. K. & Giménez-Lirola, L. G. Harness Organoid Models for Virological Studies in Animals: A Cross-Species Perspective. Front. Microbiol. 12, 725074 (2021).

25. Warburton, D. et al. The molecular basis of lung morphogenesis. Mech. Dev. 92, 55–81 (2000).

26. Zepp, J. A. & Morrisey, E. E. Cellular crosstalk in the development and regeneration of the respiratory system. Nat. Rev. Mol. Cell Biol. 20, 551–566 (2019).

27. Guimarães, C. F., Gasperini, L., Marques, A. P. & Reis, R. L. The stiffness of living tissues and its implications for tissue engineering. Nat. Rev. Mater. 5, 351–370 (2020).

28. Li, Y. et al. p63: a crucial player in epithelial stemness regulation. Oncogene 42, 3371–3384 (2023).

29. Chu, X. et al. Evidence for lung repair and regeneration in humans: key stem cells and therapeutic functions of fibroblast growth factors. Front. Med. 14, 262–272 (2020).

30. Chua, F. & Laurent, G. J. FIBROBLASTS. in Encyclopedia of Respiratory Medicine (eds. Laurent, G. J. & Shapiro, S. D.) 213–219 (Academic Press, Oxford, 2006). doi:10.1016/B0-12-370879-6/00156-3.

31. Gotoh Noriko. Control of Stemness by Fibroblast Growth Factor Signaling in Stem Cells and Cancer Stem Cells. Current Stem Cell Research & Therapy (2009).

32. Benedetto, R. et al. Epithelial Chloride Transport by CFTR Requires TMEM16A. Sci. Rep. 7, 12397 (2017).

33. Bebok, Z. et al. Failure of cAMP agonists to activate rescued ΔF508 CFTR in CFBE41o ^-^ airway epithelial monolayers: ΔF508 CFTR activity in CFBE41o ^−^ cells. J. Physiol. 569, 601–615 (2005).

34. Wang, Y., Chen, S., Yan, Z. & Pei, M. A prospect of cell immortalization combined with matrix microenvironmental optimization strategy for tissue engineering and regeneration. Cell Biosci. 9, 7 (2019).

35. Maddaly Ravi. Enhancing the Utility of Cancer Cell Lines by culturing them as 3D Aggregates and 3D Reverts – My experiences thus far. (2018) doi:10.13140/RG.2.2.33998.51521.

36. Obinata, M. The immortalized cell lines with differentiation potentials: Their establishment and possible application. Cancer Sci. 98, 275–283 (2007).

37. Soleimani, M. & Ghasemi, N. Lithium Chloride can Induce Differentiation of Human Immortalized RenVm Cells into Dopaminergic Neurons. 9, (2017).

38. Okamoto, T. et al. Clonal heterogeneity in differentiation potential of immortalized human mesenchymal stem cells. Biochem. Biophys. Res. Commun. 295, 354–361 (2002).

39. Travaglini, K. J. et al. A molecular cell atlas of the human lung from single-cell RNA sequencing. Nature 587, 619–625 (2020).

40. Basil, M. C. et al. Human distal airways contain a multipotent secretory cell that can regenerate alveoli. Nature 604, 120–126 (2022).

41. Leach, T. et al. Development of a novel air–liquid interface airway tissue equivalent model for in vitro respiratory modeling studies. Sci. Rep. 13, 10137 (2023).

42. Gontan, C. et al. Sox2 is important for two crucial processes in lung development: Branching morphogenesis and epithelial cell differentiation. Dev. Biol. 317, 296–309 (2008).

43. Laurenzi MA, Arcuri C, Rossi R, Marconi P, Bocchini V. Effects of microenvironment on morphology and function of the microglial cell line BV-2. Neurochem Res. (2001).

44. Rackley, C. R. & Stripp, B. R. Building and maintaining the epithelium of the lung. J. Clin. Invest. 122, 2724–2730 (2012).

45. Lange, K. Fundamental role of microvilli in the main functions of differentiated cells: Outline of an universal regulating and signaling system at the cell periphery. J. Cell. Physiol. 226, 896–927 (2011).

46. Agraval, H., Sharma, J. R., Dholia, N. & Yadav, U. C. S. Air–Liquid Interface Culture Model to Study Lung Cancer-Associated Cellular and Molecular Changes. in Cancer Biomarkers (ed. Deep, G.) vol. 2413 133–144 (Springer US, New York, NY, 2022).

47. Yusuf S. Khan. David T. Lynch. Histology, Lung. (StatPearls, 2023).

48. Mizushima, N. Autophagy: process and function. Genes Dev. 21, 2861–2873 (2007).

49. Li, X. et al. The Role of Autophagy in Lamellar Body Formation and Surfactant Production in Type 2 Alveolar Epithelial Cells. Int. J. Biol. Sci. 18, 1107–1119 (2022).

50. Wright, J. R. Immunoregulatory functions of surfactant proteins. Nat. Rev. Immunol. 5, 58– 68 (2005).

51. Szklarczyk, D. et al. The STRING database in 2021: customizable protein–protein networks, and functional characterization of user-uploaded gene/measurement sets. Nucleic Acids Res. 49, D605–D612 (2021).

52. Brodland, G. W. How computational models can help unlock biological systems. In Seminars in cell & developmental biology vol. 47 62–73 (Elsevier, 2015).

53. The GenOMICC Investigators et al. Genetic mechanisms of critical illness in COVID-19. Nature 591, 92–98 (2021).

54. Choi, U. Y., Kang, J.-S., Hwang, Y. S. & Kim, Y.-J. Oligoadenylate synthase-like (OASL) proteins: dual functions and associations with diseases. Exp. Mol. Med. 47, e144–e144 (2015).

55. Zani, A. et al. Interferon-induced transmembrane proteins inhibit cell fusion mediated by trophoblast syncytins. J. Biol. Chem. 294, 19844–19851 (2019).

56. Hoffmann, M. et al. SARS-CoV-2 Cell Entry Depends on ACE2 and TMPRSS2 and Is Blocked by a Clinically Proven Protease Inhibitor. Cell 181, 271–280.e8 (2020).

57. Shen, L. W., Mao, H. J., Wu, Y. L., Tanaka, Y. & Zhang, W. TMPRSS2: A potential target for treatment of influenza virus and coronavirus infections. Biochimie 142, 1–10 (2017).

58. Essalmani, R. et al. Distinctive Roles of Furin and TMPRSS2 in SARS-CoV-2 Infectivity. J. Virol. 96, e00128–22 (2022).

59. Johnson, B. A., et al. Furin Cleavage Site Is Key to SARS-CoV-2 Pathogenesis. http://biorxiv.org/lookup/doi/10.1101/2020.08.26.268854 (2020) doi:10.1101/2020.08.26.268854.

60. Xia, S. et al. The role of furin cleavage site in SARS-CoV-2 spike protein-mediated membrane fusion in the presence or absence of trypsin. Signal Transduct. Target. Ther. 5, 92 (2020).

61. Davies A, Moores C. STRUCTURE OF THE RESPIRATORY SYSTEM, RELATED TO FUNCTION. The Respiratory System. in The respiratory system (2010).

62. Kia’i N, Bajaj T. Histology, Respiratory Epithelium. 2024. (In: StatPearls [Internet]. Treasure Island, 2024).

63. Hulo, C. et al. Bacterial Virus Ontology; Coordinating across Databases. Viruses 9, 126 (2017).

