## Supplementary information. for "Design, infectability, and transcriptomic analysis of transregionally differentiated and scalable lung organoids derived from adult bronchial cells"

| Organoid Type | Source of cells | Methods and exogenous factors used | Product description | Limitations | Applications |
|---|---|---|---|---|---|
| ***ASC derived organoids*** | | | | | |
| ***Alveospheres*** | Primary human Alveolar type I and type II cells[1] | Distal lung epithelial cells are mixed in Matrigel with human lung fibroblasts (MRC5).<br><br>Pneumacult ALI medium is used to culture the organoids on thin, porous membranes.<br><br>Exogenous factors are used during the entire time of the culture:<br>50 mg per ml of Gentamycin for the first 24 hr.<br>10mM Rho kinase inhibitor for the first 48 hr.<br>2 mM of the Wnt pathway activator is used during the entire time of the culture. | Express surfactant protein C.<br><br>Maturation time 11 to 18 days. | Only resembles the distal region of the lung.<br><br>Spherical morphology. | SARS-CoV-2 infection.<br><br>Drug-screening: Remdesivir, hydroxychloroquine, interferon β |
| ***Broncheospheres*** | Primary human bronchial epithelial cells[2,3] | Bronchial epithelial cells cultured in differentiation medium with Matrigel are plated in 96-well plate containing Matrigel.<br><br>Exogenous factors: 100 ng/ml of Pam3CSK4 is applied during the whole time of culture, and Notch2 to induce goblet cell differentiation. | Prescence of goblet, basal and ciliated cells.<br><br>Notch2 and IL-17A induces goblet cell differentiation.<br><br>Maturation time 30 days. | Only resembles proximal region of the lung, without secretory or other rare cell types.<br><br>Spherical morphology | Effect of bacterial flagellin and LPS. |
| ***Human Airway organoids*** | Primary human bronchial epithelial cells from deep lung tissues of patients with cancer[4]. | Bronchial epithelial cells cultured in human airway organoid medium and 'PneumaCult' medium immersed in Matrigel.<br><br>Human organoid medium containing multiple exogenous factors such as: FGF10, FGF7, Noggin, Rspondin1.<br>'Pneumacult' medium. | Prescence of ciliated basal, secretory, and goblet cells.<br><br>Higher percentage of ciliated, secretory and goblet cells when cultured in 'PneumaCult' medium.<br><br>Maturation time 19-23 days. | Only resembles proximal region of the lung.<br><br>Spherical morphology.<br><br>The 3D structure of the organoids needs to be disrupted to be infected. | Assess influenza virus strains: H1N1, H7N9, H7N2[4]<br><br>Study of cystic fibrosis, lung cancer and Respiratory syncytial virus infection[5]. |

| | | | | | |
|---|---|---|---|---|---|
| ***Vascularized human airway organoid*** | Primary epithelium, endothelium, and lung fibroblasts[6]. | Mixed cell population embedded on Matrigel and cultured with 'Pneumacult'. | Spherical model with small protuberances.<br><br>Expression of goblet, ciliated, basal, AT1 and AT2 cells.<br><br>Maturation time 21 days. | | Possible regenerative medicine applications. |
| ***Tracheospheres*** | Primary epithelium from mouse/human tracheas[7,8]. | Tracheal epithelial cells embedded in Matrigel cultured in thin, porous membrane.<br><br>Cultured in ALI interface.<br><br>MTEC/Plus medium containing Retinoic acid, Insulin, Epidermal Growth Factor, and FBS[8,9]. | Expressing basal, secretory, and ciliated cells.<br><br>Maturation time 9 days.<br><br>Organoids can be expanded at least twice. | Only resembles the proximal airways (trachea) without goblet or secretory cells. | *NA* |
| ***Biopotential organoid*** | Primary lung epithelial cells from diseased-deep tissues[10]. | To derive proximal lung organoids, airway organoids culture medium was used (Table 1.2)<br><br>To derive distal lung organoids, distal differentiation medium was used supplemented with 50 nM dexamethasone, 100 μM 8-bromo-cAMP, 100 μM IBMX, 2% B-27,<br>Wnt agonist CHIR99021.<br><br>Organoids were embedded in Matrigel in low attachment plates. | Distal lung organoids have AT1 and AT2 cells. | Spherical organoids.<br><br>The organoids represent either proximal or distal regions of the lung (no both)<br><br>The 3D structure of the organoids needs to be broken and cultured as 2D, before infection | SARS-CoV-2 Omicron Variant infection |
| ***'Complete' lung organoid*** | Primary lung epithelial cell obtained from deep lung biopsies obtained from the normal regions of lung lobes surgically resected from lung cancer tissues[11]. | Organoids were embedded in Matrigel and cultured with lung expansion medium.<br><br>Lung expansion medium containing Wnt3, Rspondin, and Noggin, supplemented with recombinant growth factors:<br>B27, TGF-β receptor inhibitor, antioxidants, p38 MAPK inhibitor, FGF 7, | Prescence of secretory, basal, goblet, ciliated cells, as well ad AT1 and AT2 cells.<br><br>Maturation time: 7-10 days.<br><br>Organoids can be expanded. | Spherical lung organoids without specific patterning.<br><br>To be infected the organoids need to be broken and culture in 2D. | SARS-CoV-2 infection. |

| | | | | | |
|---|---|---|---|---|---|
| | | FGF 10, and ROCK inhibitor. | | | |
| ***3dGRO™ Human Lung Organoids (PDXO.149-Sigma Aldrich) | Primary lung cancer cells from various lung regions (squamous, acinar, papillary, solid, mucinous)[12]. | Not available | Prescence of AT2 cells, basal, goblet, and ciliated cells, as well as pulmonary endoderm.<br><br>Expression of ACE2 and TMPRSS receptors<br><br>Maturation time 40-60 days. | Long maturation time.<br><br>Only cancer organoids available.<br><br>No expression of AT1 cells, and secretory cells. | Possibility to use them for viral infections studies. |
| ***iPSC derived organoids*** | | | | | |
| ***Lung organoid*** | iPSC[13] | iPSC differentiated into foregut were embedded in Matrigel.<br><br>Organoid culture medium was supplemented with diverse exogenous factors varying according to specific culture times. As specified in Table 1.3 some of these molecules are FGF10, FGF4, FGF7. | Fetal airway like epithelium surrounded by mesenchyme.<br><br>Presence of goblet, ciliated, basal, AT1 and AT2 cells.<br><br>Tubular and branching morphology.<br><br>Maturation time 85-185 days (considering the initial iPSC differentiation into foregut endoderm) | Lack of maturation, it represents the fetal stage of the lung.<br><br>Inability to study adult viral diseases.<br><br>Long term culture.<br><br>Required to be transplanted into mice to improve maturation. | Evaluate human fetal lung malformations[14].<br><br>Model RSV[13]. |
| ***Airway lung organoid*** | iPSC[15] | Definitive endoderm stage cells were embedded on Matrigel and cultured with medium containing FGF10, retinoic acid, Wnt suppressor, and Notch inhibitor at different times. | Resembles the proximal airways of the lung expressing secretory, ciliated, and basal cells.<br><br>Maturation time around 46 days (considering initial cell differentiation) | The organoids mimic only the proximal region of the lung.<br><br>Spherical morphology<br><br>Long maturation time | Cystic fibrosis study. |

| | | | | | |
|---|---|---|---|---|---|
| ***Proximal airway epithelial spheroids*** | iPSC[16] | Organoids cultured in medium with Noggin, BMP4. Followed by medium with Notch inhibitor to promote ciliated cells differentiation. | Prescence of ciliated, goblet, basal, and secretory cells<br><br>Maturation time 42 days-56 days (long time higher number of ciliated cells) | The organoids mimic only on the proximal region of the lung.<br><br>Spherical morphology<br><br>Long maturation time | NA |
| ***Lung organoids*** | Embryonic cells, and iPSC[17] | Stem cells were seeded on decellularized human lung matrices on 96 well plates in presence of FGF10. | Prescence of Basal, ciliated, and secretory cells, and AT2 cells in lower number, with adjacent mesenchyme.<br><br>Maturation time 110 days. | Spherical structure.<br><br>Immature phenotype | NA |

S. Table 1. Lung organoid models, characteristics, and limitations

| Proteins upregulated in LOp vs cell monolayers | | | | |
|---|---|---|---|---|
| GLUD1 | AP2S1 | IGFBP5 | MIX23 | ATP5F1A |
| PDIA4 | CORO1A | TNXB | TPD52L1 | YWHAZ |
| CLTC | GCSH | PSMB1 | MYLK | MYO1B |
| SAFB | MATN1 | UQCRB | SPCS2 | SPTBN1 |
| PSMA3 | CSNK2A2 | GSTT2B | USP10 | ANXA8 |
| VAPA | MYH1 | DCN | GALNT2 | OAT |
| SRSF1 | CLSTN1 | PTMA | RBBP4 | H3C1 |
| IDE | NOL3 | CLEC3B | ITIH3 | AP2S1 |
| CDC42 | RBBP9 | LTF | RPS26 | CORO1A |
| KRT2 | FCGBP | FGG | RPS15 | TIMP3 |
| AP2B1 | NDRG2 | FGA | RPS14 | RPA1 |
| SF1 | XPO7 | MB | UFM1 | ABI3BP |
| TM9SF3 | NIPSNAP3A | HBD | CNN1 | HIKESHI |
| VRK1 | BCCIP | PEX14 | PRELP | ACTN4 |
| CPNE1 | ECHDC1 | SH3BGRL | GATM | MCM3 |
| MICALL1 | CACYBP | CPNE3 | TFPI2 | CD81 |
| FMOD | SFXN3 | CHAD | ECI1 | MCAM |
| DST | RBM42 | PODXL | COL15A1 | ISCA2 |
| LYZ | NTPCR | CCN1 | NUP62 | H1-1 |
| SEPHS1 | SRPK1 | TSC22D4 | GGT5 | |
| L1CAM | TP53RK | TMED7 | PPM1A | |
| SMARCA1 | FAM3D | MCTS1 | GPC1 | |
| AZGP1 | YIPF5 | FBXO2 | ATP5F1D | |
| CRYAB | ABI1 | RAB1B | CD82 | |
| IGLC2 | RHBDD2 | HDHD2 | RRM1 | |
| ARFGEF2 | RABL3 | DYNLL2 | CES1 | |
| SLC12A7 | C5orf22 | NCSTN | BGN | |
| POMP | AOC3 | RRP12 | OGN | |
| FKBP11 | SYPL1 | LAMA3 | FST | |
| THUMPD1 | LAMA4 | CRKL | ACAN | |
| SMC4 | CST6 | MKI67 | PIP | |
| RBKS | PEA15 | MCM5 | SPARC | |
| ANP32E | KRT81 | VCAN | RBP1 | |
| GDF15 | GOLGA4 | CA1 | ISG15 | |
| GALM | SRSF5 | PLG | FGB | |
| CEMIP | STK4 | QSOX1 | KRT14 | |
| SETD3 | COL14A1 | PPP6C | HLA-DRA | |
| CCDC80 | AKAP12 | TM9SF4 | PLAU | |
| CDC73 | ALDH6A1 | TIMM9 | DDX39A | |
| PDCD4 | PLCB3 | LSM7 | ARL6IP5 | |
| ATP6AP1 | SNRPG | ATG3 | KPNA3 | |
| TPSAB1 | TMEM33 | CTPS2 | ETF1 | |
| PABPC4 | OXCT1 | NUP88 | DLAT | |
| CBX3 | LAMB2 | PPP1R14B | KPNB1 | |
| FSTL1 | PLTP | COPS5 | TXN | |
| KIF23 | LUM | HTRA1 | MYOF | |
| DYNLL1 | SSR4 | SELENOM | SLC3A2 | |
| POLR2L | UQCRFS1 | NUP210 | DNAJC9 | |

S. Table 2. Proteins upregulated in LOp in comparison to HBE and IMR-90.

| Proteins upregulated in LOi vs cell monolayers | | | | | |
|---|---|---|---|---|---|
| KRT2 | CES1 | SSR4 | CPNE1 | VAPA | STK4 |
| HSPD1 | TNXB | UQCRFS1 | GALM | CANX | DST |
| SRSF1 | MATN1 | ECI1 | FAM3D | CLTC | PLCB3 |
| KPNB1 | BGN | NUP62 | HTRA1 | GLUD1 | RPS15 |
| KPNA3 | PSMB1 | GGT5 | TM9SF4 | POR | SNRPG |
| PSMA3 | PIP | TIMP3 | NUP210 | S100A4 | LYZ |
| SORD | GSTT2B | SHMT1 | MICALL1 | SERPINB1 | PRELP |
| SAFB | KRT14 | L1CAM | ABI1 | TXN | POMP |
| IMPDH2 | CRYAB | ATP5F1D | ISCA2 | AP2B1 | NDRG2 |
| CKMT1A | CLSTN1 | RPA1 | ABI3BP | ANXA8 | MCTS1 |
| FASN | HLA-DRA | AZGP1 | TPD52L1 | EPHA2 | LSM7 |
| ACO2 | COPS5 | GCSH | CST6 | ERMP1 | XPO7 |
| RSL1D1 | RAB1B | FST | MYLK | PPID | THUMPD1 |
| ATP5F1A | HDHD2 | RBP1 | PEA15 | RPS28 | ECHDC1 |
| CDC42 | SRPK1 | CLEC3B | KRT81 | RPL31 | ATG3 |
| MYOF | YIPF5 | ISG15 | GOLGA4 | SPTBN1 | RBKS |
| DLAT | TPSAB1 | FGG | SRSF5 | ADA | ANP32E |
| ETF1 | FSTL1 | HBD | CBX3 | SERPINB5 | VRK1 |
| HSPA8 | DYNLL1 | SH3BGRL | FMOD | IDH2 | PTMA |
| YWHAZ | POLR2L | NOL3 | ITIH3 | FABP5 | FGB |
| MYO1B | RPS14 | CPNE3 | COL14A1 | TBCA | MB |
| IDE | GPC1 | TSC22D4 | AKAP12 | GPI | PLAU |
| TIMM9 | CD82 | FBXO2 | RPS26 | PFN2 | PLG |
| TMED7 | RRM1 | SFTPA1 | TMEM33 | ACTN4 | ARL6IP5 |
| NIPSNAP3A | LTF | HIKESHI | LAMB2 | BTF3 | CCN1 |
| TM9SF3 | RBBP9 | MIX23 | GATM | ALDH6A1 | CHAD |
| RBM42 | DDX39A | C5orf22 | SEPHS1 | KIF23 | SYPL1 |
| NTPCR | PPP6C | AOC3 | CRKL | CD81 | CTPS2 |
| NUP88 | BCCIP | SF1 | MKI67 | PLTP | PDIA4 |
| TP53RK | FKBP11 | UFM1 | MCAM | AP2S1 | |
| DYNLL2 | SMC4 | OXCT1 | COL15A1 | LUM | |
| NCSTN | CACYBP | IGLC2 | PPM1A | TFPI2 | |
| CCDC80 | SFXN3 | FGA | CORO1A | MCM5 | |
| CDC73 | GDF15 | CA1 | OGN | SMARCA1 | |
| RRP12 | PPP1R14B | PEX14 | CSNK2A2 | IGFBP5 | |
| PDCD4 | SELENOM | QSOX1 | ACAN | CES1 | |
| LAMA4 | CEMIP | PODXL | UQCRB | LAMA3 | |
| ATP6AP1 | SETD3 | FCGBP | VCAN | SPCS2 | |
| GALNT2 | RHBDD2 | ARFGEF2 | SPARC | USP10 | |
| RBBP4 | RABL3 | SLC12A7 | DCN | PABPC4 | |

S. Table 3. Proteins upregulated in LOi in comparison to CFBE and IMR-90.

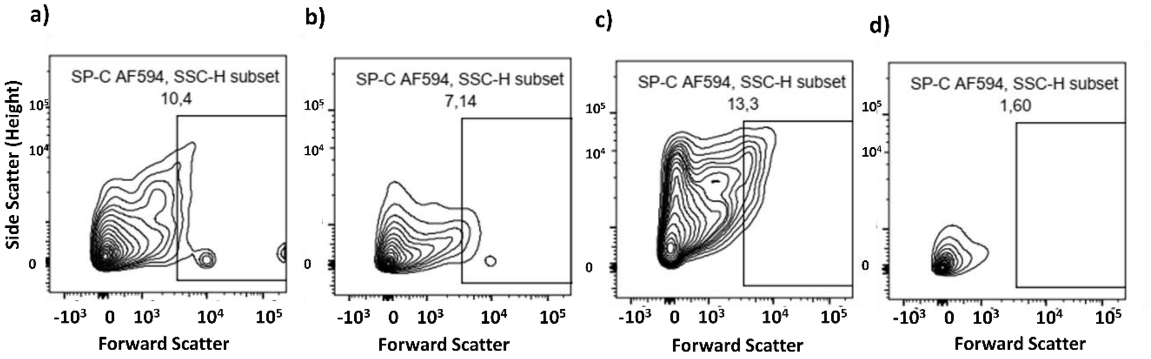


S. Fig. 1 Flow cytometry to confirm AT2 cells of organoids derived by primary cells LOp. a-c) Percentage of SFTPC+ (AF594) cells detected in each organoid. d) Fluorescence minus one (FMO) 594 was used to calibrate the instrument. N=3

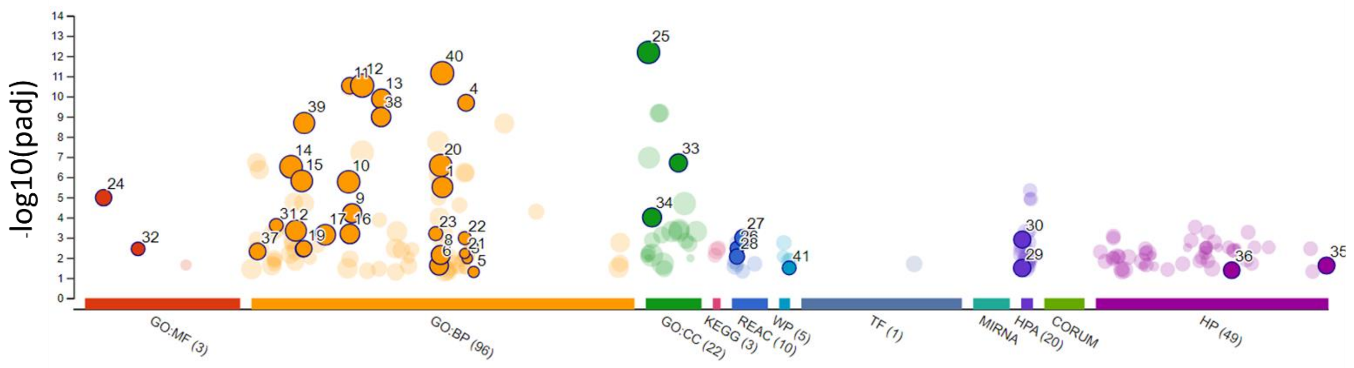


| ID | Term ID | Term name |
|---|---|---|
| 1 | GO:0048870 | cell motility |
| 2 | GO:0008283 | cell population proliferation |
| 3 | GO:0060638 | mesenchymal-epithelial cell signaling |
| 4 | GO:0060541 | respiratory system development |
| 5 | GO:0061140 | lung secretory cell differentiation |
| 6 | GO:0048598 | embryonic morphogenesis |
| 7 | GO:0048754 | branching morphogenesis of an epithelial tube |
| 8 | GO:0048729 | tissue morphogenesis |
| 9 | GO:0030855 | epithelial cell differentiation |
| 10 | GO:0030154 | cell differentiation |
| 11 | GO:0030324 | lung development |
| 12 | GO:0032502 | developmental process |
| 13 | GO:0035295 | tube development |
| 14 | GO:0007275 | multicellular organism development |
| 15 | GO:0009653 | anatomical structure morphogenesis |
| 16 | GO:0030334 | regulation of cell migration |
| 17 | GO:0016477 | cell migration |
| 18 | GO:0009799 | specification of symmetry |
| 19 | GO:0009855 | determination of bilateral symmetry |
| 20 | GO:0048731 | system development |
| 21 | GO:0060435 | bronchiole development |
| 22 | GO:0060441 | epithelial tube branching involved in lung morphogenesis |
| 23 | GO:0048286 | lung alveolus development |
| 24 | GO:0005201 | extracellular matrix constituent |
| 25 | GO:0005576 | extracellular region |
| 26 | GO:0030312 | collagen chain trimerization |
| 27 | REAC:R-HSA-8948216 | extracellular matrix organization |
| 28 | REAC:R-HSA-1474244 | collagen formation |
| 29 | REAC:R-HSA-1474290 | bronchus, ciliated cells (cilia axoneme) |
| 30 | HPA:0060862 | bronchus, ciliated cells (tip of cilia) |
| 31 | HPA:0060883 | epithelium cilia movement involved in extracellular fluid movement |
| 32 | GO:0003351 | extracellular matrix structural constituent conferring tensile strength |
| 33 | GO:0030020 | collagen-containing extracellular matrix |
| 34 | GO:0062023 | cilium |
| 35 | GO:0005929 | Abnormal lung development |
| 36 | HP:4000059 | Abnormal bronchus morphology |
| 37 | HP:0025426 | morphogenesis of a branching structure |
| 38 | GO:0001763 | tube morphogenesis |
| 39 | GO:0035239 | tissue development |
| 40 | GO:0009888 | anatomical structure development |
| 41 | GO:0048856 | miRNA targets in ECM and membrane receptors |

S. Fig. 2 Functional Enrichment Analysis. A Functional Enrichment analysis of genes with major change between cell monolayers and organoids (p adj.<0.01) by G: Profiler. Abbreviations: MF-Molecular Function, BP: Biological Process. CC-Cellular Component. KEGG- Kyoto Encyclopedia of Genes and Genomes. REAC - Reactome pathways. WP – WikiPathways. TF - Transfac transcription factor binding site predictions. MIRNA - mirTarBase miRNA targets. HPA - Human Protein Atlas expression data. CORUM - Manually annotated protein complexes from mammalian organisms. HP - Human Phenotype Ontology, a standardized vocabulary of phenotypic abnormalities encountered in human disease.

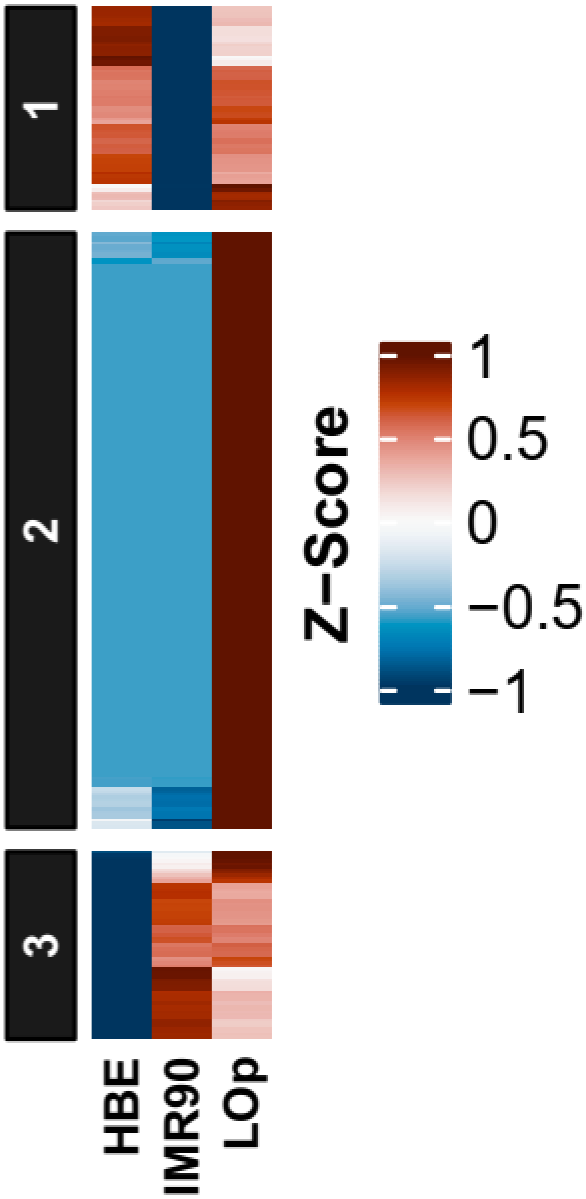


S.Fig. 3 Proteomic profile of organoids derived from primary cells, LOp, and its comparison with HBE and IMR-90 cell monolayers. Cluster 1 shows the similarities between HBE and LOp. Cluster 2 highlights the proteins upregulated in the organoid in respect to cell monolayers. Cluster 3 shows the similarities of the proteome between IMR-90 and LOp. Grouped by abundances, linked by average. N=3. Upregulation shown in red. Downregulation shown in blue.

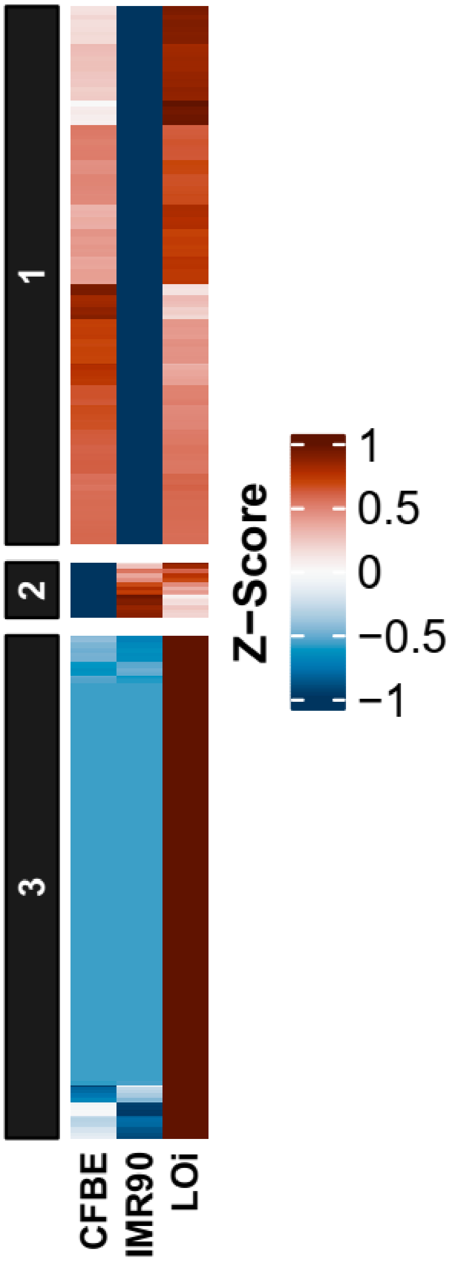


S.Fig. 4 Proteomic profile of organoids derived from immortalized cells, LOi, and its comparison with CFBE and IMR-90 cell monolayers. Cluster 1 shows the similarities between CFBE and LOi. Cluster 2 shows the similarities of the proteome between IMR-90 and LOi. Cluster 3 highlights the proteins upregulated in the organoids, LOi, in respect to cell monolayers. Grouped by abundances, linked by average. N=3. Upregulation shown in red. Downregulation shown in blue.

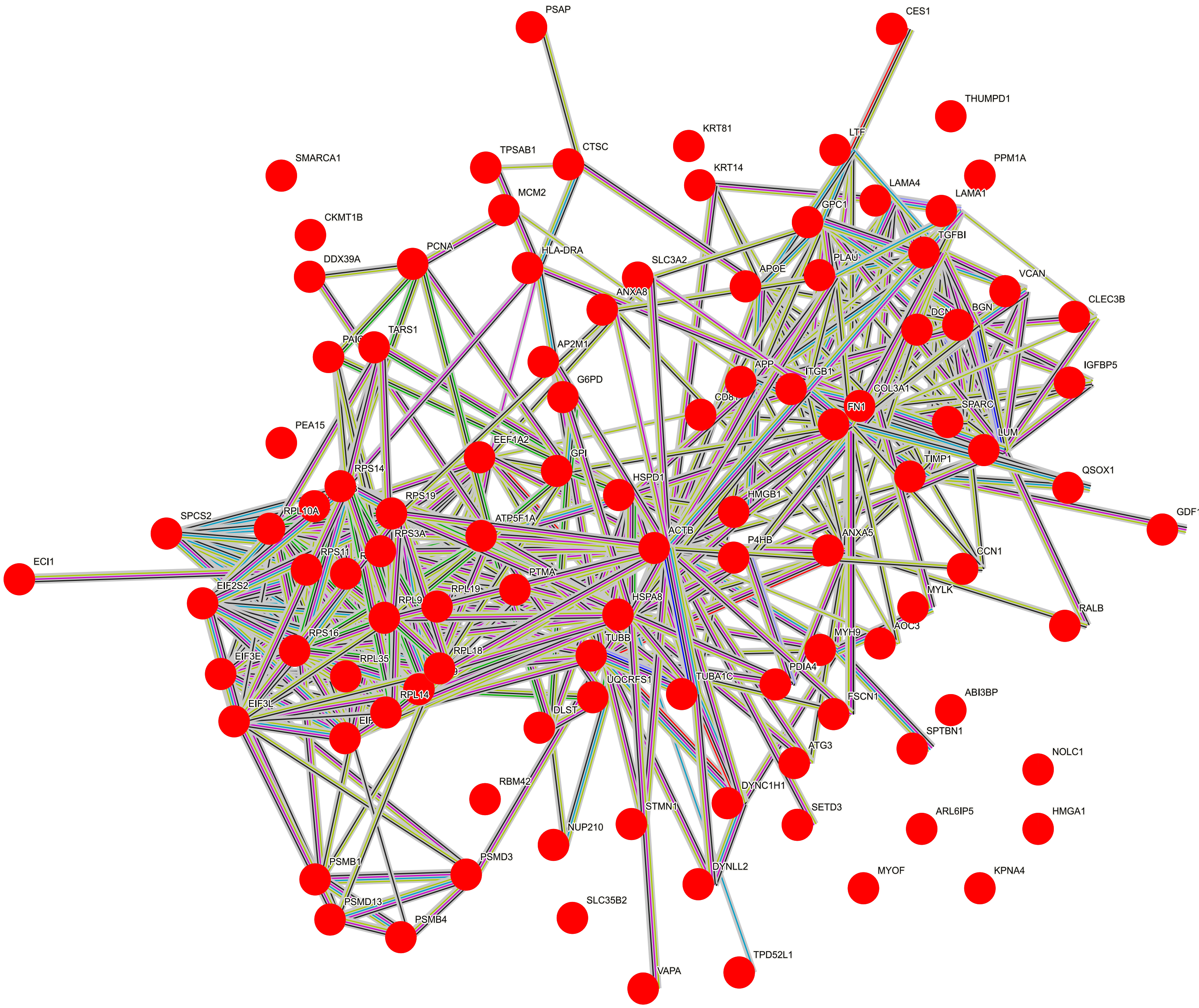


S. Fig. 5 Upregulated proteins in LOp with high specificity to the lung showing connections on pathways. Generated by STRING. Pathways colors: Blue-from curated database, Pink-\experimentally determined. Green-Gene neighborhood. Red-Gene fusions. Blue-Gene co-occurrence. Yellow-Textmining. Black co-expression. Dark blue: protein homology.

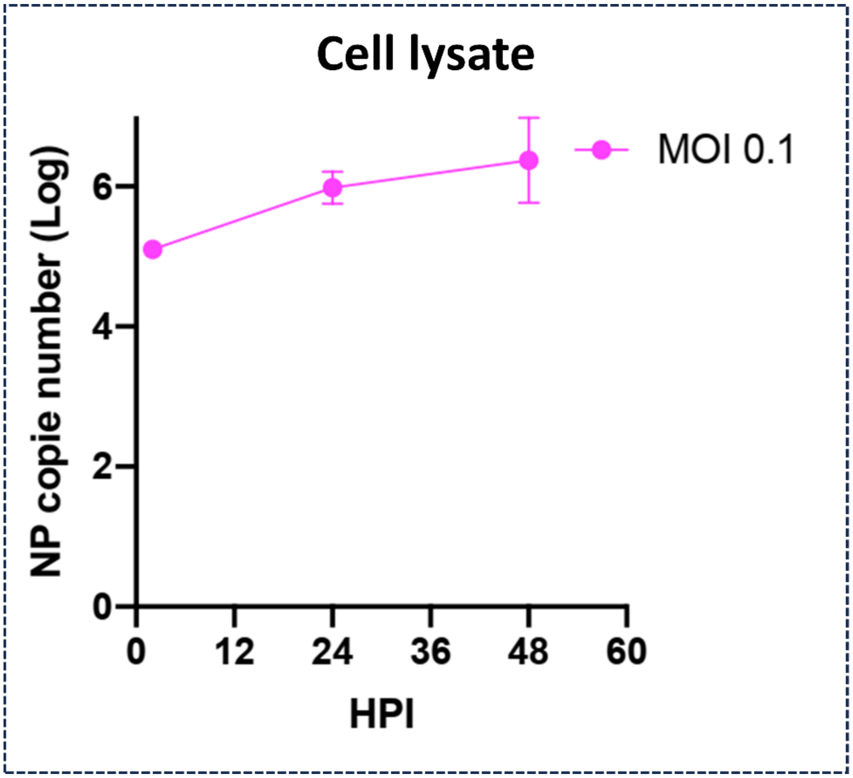


S.Fig.6 Nucleocapsid protein load (log) in organoid lysate after 24 hours post infection, and 48 hours post infection of influenza H1N1-PR-8. N=3

### References

1. Mulay, A. *et al.* SARS-CoV-2 infection of primary human lung epithelium for COVID-19 modeling and drug discovery. *Cell reports* **35**, 109055 (2021).
2. Sprott, R. F. *et al.* Flagellin shifts 3D bronchospheres towards mucus hyperproduction. *Respir Res* **21**, 222 (2020).
3. Danahay, H. *et al.* Notch2 Is Required for Inflammatory Cytokine-Driven Goblet Cell Metaplasia in the Lung. *Cell Reports* **10**, 239–252 (2015).
4. Zhou, J. *et al.* Differentiated human airway organoids to assess infectivity of emerging influenza virus. *Proc. Natl. Acad. Sci. U.S.A.* **115**, 6822–6827 (2018).

5. Sachs, N. *et al.* Long-term expanding human airway organoids for disease modeling. *The EMBO Journal* **38**, e100300 (2019).

6. Tan, Q., Choi, K. M., Sicard, D. & Tschumperlin, D. J. Human airway organoid engineering as a step toward lung regeneration and disease modeling. *Biomaterials* **113**, 118–132 (2017).

7. Cunniff, B., Druso, J. E. & van der Velden, J. L. Lung organoids: advances in generation and 3D-visualization. *Histochem Cell Biol* **155**, 301–308 (2021).

8. Rock, J. R. *et al.* Basal cells as stem cells of the mouse trachea and human airway epithelium. *Proc Natl Acad Sci USA* **106**, 12771 (2009).

9. *Epithelial Cell Culture Protocols: Second Edition*. vol. 945 (Humana Press, Totowa, NJ, 2013).

10. Chiu, M. C. *et al.* A bipotential organoid model of respiratory epithelium recapitulates high infectivity of SARS-CoV-2 Omicron variant. *Cell Discov* **8**, 57 (2022).

11. Tindle, C. *et al.* Adult stem cell-derived complete lung organoid models emulate lung disease in COVID-19. *eLife* **10**, e66417 (2021).

12. Millipore Sigma. 3dGRO$^{TM}$ Human Lung Organoids (PDXO.149). https://www.sigmaaldrich.com/CA/en/product/mm/scc607#product-documentation.

13. Miller, A. J. *et al.* Generation of lung organoids from human pluripotent stem cells in vitro. *Nat Protoc* **14**, 518–540 (2019).

14. Chen, Y.-W. *et al.* A three-dimensional model of human lung development and disease from pluripotent stem cells. *Nat Cell Biol* **19**, 542–549 (2017).

15. McCauley, K. B. *et al.* Efficient Derivation of Functional Human Airway Epithelium from Pluripotent Stem Cells via Temporal Regulation of Wnt Signaling. *Cell Stem Cell* **20**, 844-857.e6 (2017).

16. Konishi, S. *et al.* Directed Induction of Functional Multi-ciliated Cells in Proximal Airway Epithelial Spheroids from Human Pluripotent Stem Cells. *Stem Cell Reports* **6**, 18–25 (2016).

17. Dye, B. R. *et al.* In vitro generation of human pluripotent stem cell derived lung organoids. *eLife* **4**, e05098 (2015).
